# Metabolic-secretory decoupling defines a disease-intrinsic state in rheumatoid arthritis monocytes

**DOI:** 10.64898/2026.04.23.720351

**Authors:** Shao Thing Teoh, Silja Malkewitz, Cristian Iperi, Celia Makowiec, Asimina Kakale, Seyram Maureen Duphey, Anastasiya Börsch, Katarzyna Buczak, Witold Wolski, Ming Yang, Christian Frezza, Caroline Ospelt, Oliver Distler, Diego Kyburz, Bojana Mueller-Durovic

**Affiliations:** Center for Experimental Rheumatology, University of Zurich and University Hospital of Zurich, Department of Rheumatology, Zurich-Schlieren, Switzerland; Immunobiology Laboratory, Department of Biomedicine, University of Basel and University Hospital of Basel, Basel, Switzerland; Bioinformatics Core Facility, Department of Biomedicine, University of Basel, Basel, Switzerland; Proteomics Core Facility, Biozentrum, University of Basel, Basel, Switzerland; Proteomics Unit, Functional Genomics Center Zurich, ETH Zürich and the University of Zurich, Zurich, Switzerland; Institute for Metabolomics in Ageing, Faculty of Medicine and University Hospital Cologne, Cluster of Excellence Cellular Stress Responses in Aging-associated Diseases (CECAD), University of Cologne, Cologne, Germany; Institute of Genetics, Faculty of Mathematics and Natural Sciences, Cluster of Excellence Cellular Stress Responses in Aging-associated Diseases (CECAD), University of Cologne, Cologne, Germany; Center for Molecular Medicine (CMMC), University of Cologne, Cologne, Germany; Experimental Rheumatology Laboratory, Department of Biomedicine, University of Basel, Basel, Switzerland

## Abstract

**Objectives:** Circulating monocytes from rheumatoid arthritis (RA) patients are pre-primed for inflammatory activation, but their disease-intrinsic features have not been systematically characterized. Given the important role of metabolism in shaping immune cell function, we aimed to determine how this pre-primed state is underpinned metabolically and whether these changes persist across different activation states, using an unbiased multi-omics approach.

**Methods:** Peripheral blood CD14⁺ monocytes from RA patients and matched healthy donors were analyzed in an undifferentiated state (M0) and after differentiation into classically activated M(IFNγ+LPS) and alternatively activated M(IL-4) macrophages, followed by acute lipopolysaccharide (LPS) stimulation. Metabolomic (untargeted LC–MS/MS), transcriptomic (RNA-seq), and proteomic (label-free mass LC-MS/MS) profiling were performed. Data was comprehensively analyzed by weighted gene correlation network analysis, differential analysis, gene set enrichment analysis, multi-omics factor analysis and metabolic flux modeling.

**Results:** RA monocytes exhibited a stable disease-driven signature across activation states. Integration of metabolomic, transcriptomic and proteomic data revealed an unexpected convergence on metabolic–secretory coupling, with depletion of nucleotide and redox metabolites, downregulation of mitochondrial and translational pathways, and remodeling of the secretory apparatus, including loss of cis-Golgi components. Consistently, metabolic modeling predicted reduced glycosylation fluxes, connecting metabolic changes to altered secretory capacity.

**Conclusions:** RA monocytes adopt a stable, disease-intrinsic state that persists across activation conditions. Multi-omics data identify a linked metabolic and secretory defect, with reduced glycosylation capacity as a potential functional consequence. This metabolic-secretory coupling represents a defining feature of RA monocyte dysfunction and a potential therapeutic target.

## INTRODUCTION

Rheumatoid arthritis (RA) is a chronic systemic autoimmune disease characterized by persistent synovial inflammation, progressive joint destruction, and substantial systemic manifestations. Although adaptive immune responses involving autoreactive T and B cells are central to disease initiation, innate immune cells – particularly monocytes and macrophages – are critical effectors of tissue damage and drivers of chronic inflammation (1–3). Synovial macrophage abundance correlates strongly with disease activity and radiographic progression, and depletion of synovial macrophages parallels clinical improvement following effective therapy (3, 4). Circulating monocytes serve as an important source of these tissue macrophages and therefore represent both a biologically relevant and experimentally accessible cell population for interrogating disease-associated immune dysregulation in RA.

Peripheral blood monocytes from RA patients exhibit altered activation states, enhanced migratory capacity, and dysregulated cytokine responses compared to healthy donors (5, 6). These differences likely reflect a combination of chronic inflammatory exposure and disease-intrinsic reprogramming that persists outside the joint microenvironment. Upon entering tissues, monocytes differentiate into macrophages with diverse functional phenotypes. Although macrophage activation states exist along a continuum in vivo, the binary in vitro paradigm of classically activated (M1-like) versus alternatively activated (M2-like) polarization remain widely used as a controlled system to examine how monocytes respond to defined inflammatory or anti-inflammatory cues (7, 8). **However, it remains unclear which disease-intrinsic features of RA monocytes are preserved across these activation states.**

While transcriptomic studies of monocytes and macrophages have identified inflammatory, interferon-associated, and metabolic gene expression signatures associated with disease activity and treatment response (6, 9, 10), **integrated multi-omics studies focusing specifically on primary human monocytes across controlled differentiation and inflammatory conditions remain scarce**. Integrated multi-omics approaches have demonstrated value in resolving complex immune phenotypes by identifying coordinated molecular programs that are not apparent from single-omics analyses alone (11–13). However, existing RA omics studies typically (i) focus on a single molecular layer, (ii) analyze heterogeneous cell populations or tissue samples, or (iii) assess RA-associated differences under a single activation state. For example, proteomic analyses of inflamed synovium in RA have revealed alterations in antigen presentation machinery, cytoskeletal organization, and extracellular matrix–related proteins (14–16). In parallel, metabolomics studies – predominantly performed in serum, plasma, or synovial fluid – have demonstrated profound perturbations in amino acid metabolism, lipid metabolism, redox homeostasis, and central carbon metabolism in RA(17–24). While these studies have substantially advanced understanding of RA pathogenesis, it remains unclear to what extent molecular alterations in RA monocytes are intrinsic to the disease state versus secondary to differentiation or activation programs, and whether discrepancies between different omics layers reflect technical limitations or biologically meaningful regulation.

In this study, we performed integrated transcriptomic, proteomic, and metabolomic profiling of peripheral blood monocytes and monocyte-derived macrophages from patients with RA and matched healthy donors across controlled differentiation and inflammatory conditions. This approach allowed us to characterize how disease status interacts with macrophage differentiation and inflammatory activation. By integrating gene expression, protein levels, and metabolite abundances, we provide a comprehensive framework for understanding disease-associated immune-metabolic dysregulation in RA monocytes and identify RA-associated molecular processes that may be overlooked by single-omics approaches.

## RESULTS

### Rheumatoid arthritis monocytes display a persistent metabolic signature across activation states

To define disease-associated metabolite changes in circulating monocytes and monocyte-derived macrophages from RA patients, we performed untargeted LC-MS/MS metabolomics in 6 RA patients and 6 healthy donors (HDs). Monocyte-derived macrophages classically activated (M(IFNγ+LPS)) or alternatively activated (M(IL-4)) polarization states, or with acute lipopolysaccharide (LPS) stimulation are widely used in experimental macrophage studies, yet the impact of these stimuli on RA-associated transcriptional signatures are not fully understood. We therefore analyzed RA and HD monocytes in the undifferentiated state (M0) or following in vitro differentiation into M(IFNγ+LPS) or M(IL-4) macrophages, with further acute lipopolysaccharide (LPS) stimulation (100 ng/mL for 24 h) or vehicle control prior to harvesting.

Across all samples, peak areas of 202 metabolites were obtained following untargeted LC-MS/MS data processing (peak picking, alignment, adduct and isotopes grouping, annotation). Hierarchical clustering across all conditions showed that RA samples segregated from healthy donors independent of differentiation or acute LPS stimulation (**Fig. S1A**). Most metabolites differed consistently between RA and healthy donors, indicating a strong disease-driven signature. Variance partitioning confirmed RA disease status as the dominant source of variation across the dataset (**Fig. S1B**). In healthy donors, samples were primarily structured by differentiation state (M0, M(IFNγ+LPS), M(IL-4)) and further separated by acute LPS stimulation, particularly in M0 cells. In contrast, this organization was markedly attenuated in RA, where the contribution of differentiation state to overall variance was substantially reduced (**Fig. S1C**).

Weighted gene correlation network analysis (WGCNA) of the metabolomics data identified eight modules of co-varying metabolites. The largest modules (‘blue’, ‘grey’, and ‘turquoise’) accounted for the majority of metabolites and showed strong associations with RA status (**Fig. 1A**). The ‘blue’ and ‘grey’ modules were enriched in RA, whereas the ‘turquoise’ module was depleted, indicating a broad disease-associated shift in metabolite abundance. In contrast, only a small module (‘yellow’, four metabolites) was strongly associated with M(IL-4) differentiation (r = 0.76), while differentiation and acute LPS stimulation showed comparatively limited influence on overall module structure.

**Figure 1.**
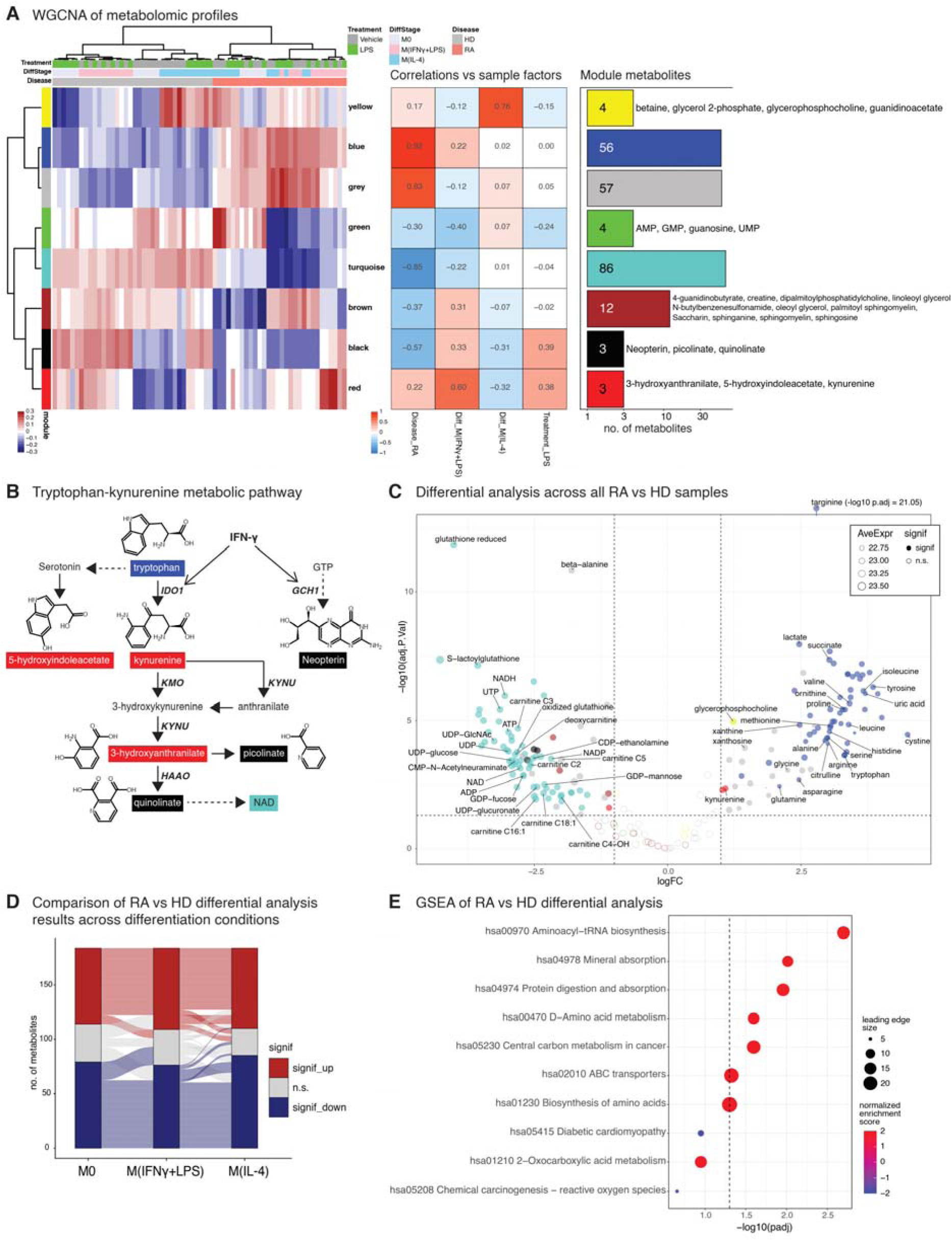
Rheumatoid arthritis monocytes display a persistent metabolic signature across activation states. **(A)** Weighted correlation network analysis (WGCNA) of untargeted metabolomics data (202 metabolites) identified eight metabolite modules. Left: clustered heatmap showing module eigengene values in individual samples, as well as clustering patterns of samples according to eigengenes. Center: heatmap of correlations between module eigengenes and sample traits including disease status (RA, HD), differentiation state (M0, M(IFNγ+LPS), M(IL-4), and acute LPS treatment. Right: bar plot indicating the number of metabolites per module. ‘Blue’ and ‘grey’ modules positively correlated with RA; ‘turquoise’ negatively correlated with RA; ‘yellow’ strongly correlated with M(IL-4). **(B)** Schematic representation of the tryptophan-kynurenine metabolic pathway. Metabolites are color coded by WGCNA module: the red module contains proximal intermediates (kynurenine, 3-hydroxyanthranilate) as well as 5-hydroxyindoleacetate (another derivative of tryptophan via the serotonin pathway), while the black module includes distal metabolites (picolinate, quinolinate) as well as Neopterin, consistent with IFN-associated metabolic responses. **(C)** Volcano plot showing differential metabolite abundance between RA and HD samples across all differentiation states. The analysis included 6 RA patients and 6 healthy donors, each profiled across 3 differentiation states and 2 treatment conditions (36 samples per group). Differential metabolite abundance was assessed using a linear mixed-effects model including donor as a random effect. The horizontal axis indicates log2 fold change, while vertical axis indicates adjusted p-value of differences. Average metabolite peak areas are indicated by data point sizes. Significant metabolites (adjusted p-value < 0.05, absolute log2FC > 1) are represented by filled data points while non-significant metabolites are represented by empty circles. Data points are color-coded according to WGCNA module. Specific metabolites are labeled to aid interpretation: RA samples show decreased nucleotides, nucleotide sugars, NAD(H)/NADP(H), glutathione-related metabolites, and carnitines, and increased amino acids and urea cycle intermediates. **(D)** Alluvial plot visualizing the persistence of the RA metabolic signature irrespective of differentiation states. Each left-to-right ‘flow’ tracks a metabolite’s RA vs HD differential abundance (significantly up-, downregulated, or not significant) across M0, M(IFNγ+LPS), and M(IL-4) subsets. **(E)** Gene Set Enrichment Analysis (GSEA) performed using the t statistic of RA vs HD differential metabolite abundance analysis. Point colors indicate normalized enrichment score (NES) whereby positive and negative scores indicate up- and downregulation in RA respectively, while point sizes indicate number of leading edge genes. The vertical dashed line indicates the adjusted p-value = 0.05 threshold. Top 10 metabolite pathways with smallest adjusted p-value are shown. Amino acid–related pathways are particularly enriched, including Aminoacyl-tRNA biosynthesis, Mineral absorption, and Protein digestion and absorption.

Notably, the ‘black’ and ‘red’ modules were enriched for metabolites of the tryptophan–kynurenine pathway (**Fig. 1B**). The ‘red’ module contained proximal intermediates (tryptophan, kynurenine, 3-hydroxyanthranilate), whereas the ‘black’ module comprised distal metabolites (picolinate and quinolinate) together with neopterin, a marker of interferon (IFN)-driven inflammatory activation. These findings suggest that the ‘black’ module captures metabolic changes associated with IFN responses. Overall, the segregation of metabolites into distinct modules linked to disease status, differentiation state, and inflammatory activation supports the biological relevance of the WGCNA-derived network structure.

Overrepresentation analysis (ORA) of the RA-associated modules showed that the ‘blue’ module was significantly enriched for amino acid-related pathways, including aminoacyl-tRNA biosynthesis, consistent with the broad increase in amino acids observed in RA monocytes (**Fig. S1D**). In contrast, the ‘grey’ and ‘turquoise’ modules showed only modest enrichment patterns that did not remain significant after multiple-testing correction, with trends toward pathways linked to central carbon metabolism and redox biology (**Fig. S1E-F**). These findings indicate that amino acid metabolism represents the most robust pathway-level feature of the RA-associated metabolic signature, whereas other metabolic alterations are more subtle.

Given the consistent RA-HD separation across conditions, we performed differential analysis across all samples. RA monocytes and macrophages showed reduced levels of nucleotides, nucleotide sugars, redox metabolites (including NAD and glutathione), and carnitines, alongside increased urea cycle intermediates (arginine, ornithine, citrulline) and many other amino acids (**Fig. 1C**). These changes were largely consistent across M0, M(IFNγ+LPS), and M(IL-4) states (**Fig. 1D**), indicating a stable disease-associated metabolic signature. Reduced nucleotide levels were not associated with patient methotrexate or leflunomide treatment status, which could potentially affect nucleotide levels (**Fig. S1G**) and were confirmed in an independent cohort utilizing monocytes immediately following CD14+ isolation (**Fig. S1H–I**). Together, these findings demonstrate a robust and persistent metabolic alteration in circulating RA monocytes, independent of differentiation or in vitro handling.

Gene set enrichment analysis (GSEA) of the RA–HD differential signature confirmed enrichment of amino acid–related pathways, consistent with increased amino acid abundance in RA samples (**Fig. 1E**) and in line with ORA results from the ‘blue’ module (**Fig. S1D**). Together with network analysis, these findings indicate that RA-associated metabolic changes are dominated by shifts in amino acid metabolism, with additional modules capturing distinct axes such as kynurenine pathway/IFN responses and alternative macrophage differentiation. Overall, disease status emerged as the primary driver of metabolic organization, exceeding the effects of differentiation state.

### Transcriptome analysis identifies rheumatoid arthritis-associated inflammatory activation and translational constraints

Transcriptomic profiling across all conditions showed that samples were primarily structured by differentiation state, with clear separation of M0, M(IFNγ+LPS), and M(IL-4) cells (**Fig. S2A**).

Acute LPS stimulation further stratified M0 and M(IL-4) samples, whereas this effect was less pronounced in M(IFNγ+LPS) macrophages, reflecting their prior LPS exposure during differentiation. Against this background, RA samples exhibited a superimposed disease-associated signature, indicating that inflammatory activation occurs within, but does not override, differentiation-defined transcriptional states.

Variance partitioning confirmed differentiation state as the dominant driver of transcriptomic variation, accounting for a median of ∼35% of total variance (**Fig. S2B**). Within individual states, the contribution of other variables differed, with acute LPS stimulation dominating in M0, donor identity in M(IFNγ+LPS), and both factors contributing in M(IL-4) (**Fig. S2C**). In contrast, disease status explained only a small fraction of variance and was confined to a limited set of genes, indicating that RA does not induce global transcriptomic rewiring but rather a focused transcriptional program. WGCNA of the transcriptomic dataset identified 22 gene modules (**Fig. 2A**). All but two modules showed an absolute correlation of at least 0.4 with one or more sample traits, including RA status, differentiation state, or LPS treatment, indicating that these variables collectively explained most module expression patterns (**Fig. 2A, middle**).

**Figure 2.**
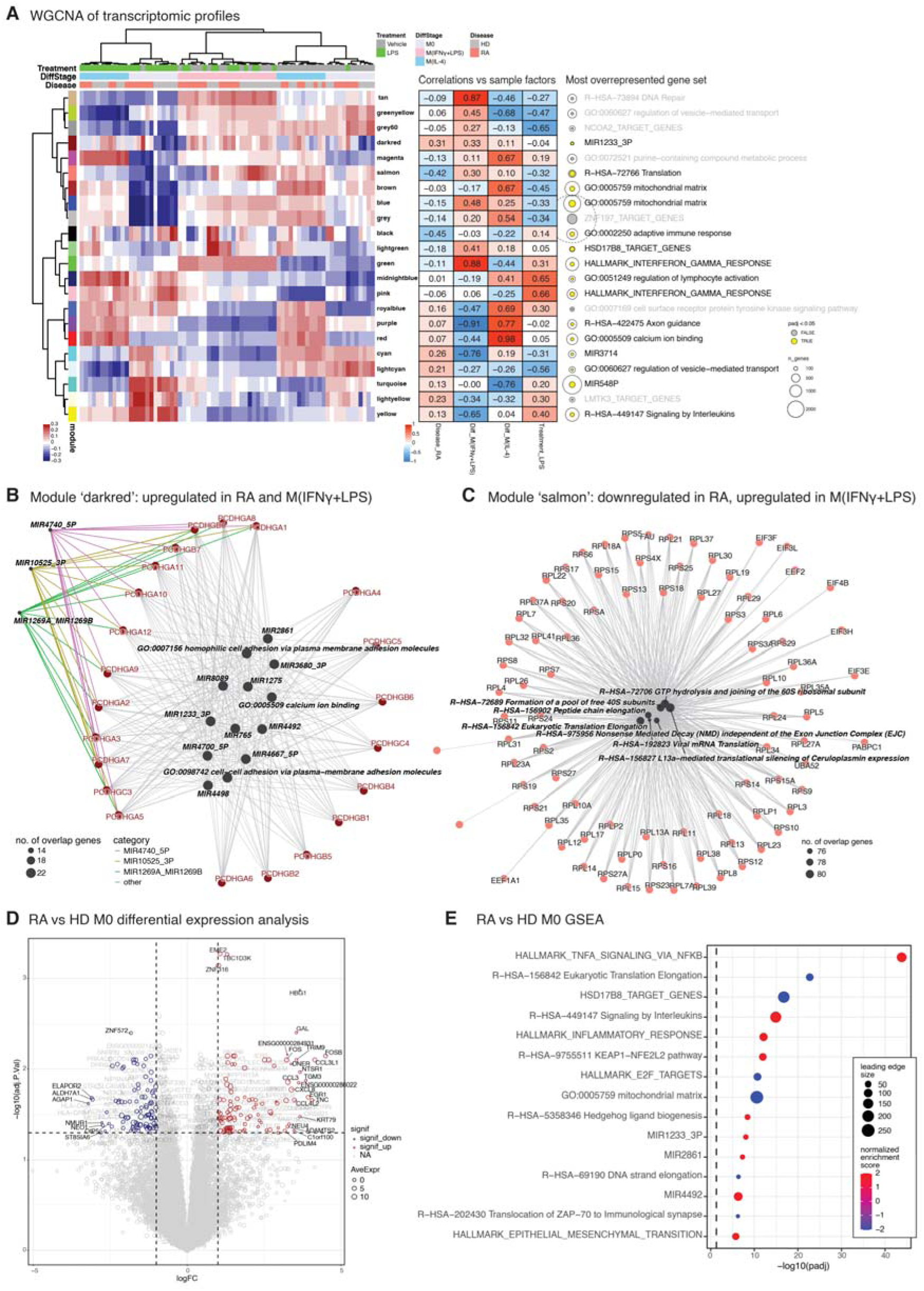
Transcriptome analysis identifies rheumatoid arthritis-associated inflammatory activation and translational constraints. **(A)** Weighted correlation network analysis (WGCNA) of transcriptomics data identified 22 gene modules. Left: clustered heatmap showing module eigengene values in individual samples, as well as clustering patterns of samples according to eigengenes. Center: heatmap of correlations between module eigengenes and sample traits including disease status (RA, HD), differentiation state (M0, M(IFNγ+LPS), M(IL-4), and acute LPS treatment. Right: top overrepresented gene set, defined as having the largest number of overlap genes, for each module. The double concentric circles next to gene set names represent number of genes in the module (size of outer circle) and number of overlap genes (size of inner circle), respectively. The inner circle is color coded by statistical significance of the top overrepresented gene set (yellow if adjusted p-value < 0.05, grey otherwise). Modules ‘black’, ‘salmon’, and ‘darkred’ show moderate correlation with RA. **(B)** Network representation of overlap between gene sets significantly enriched within the ‘darkred’ module. Gene sets are represented by dark grey filled circles with sizes according to number of overlap genes. Module genes are represented by dark red points. Multiple predicted miRNA target gene sets including *MIR1233_3P* are enriched within this module, with enrichment driven predominantly by clustered protocadherin genes (*PCDHGAx*/*PCDHGBx*). **(C)** Network representation of overlap between gene sets significantly enriched within the ‘salmon’ module. Gene sets are represented by dark grey filled circles with sizes according to number of overlap genes. Module genes are represented by salmon-colored points. Gene set enrichment within this module is primarily driven by ribosomal proteins and translation initiation/elongation factors. **(D)** Volcano plot showing RA vs HD differential gene expression in untreated M0 monocytes. The analysis included 6 RA patients vs 6 healthy donors. The horizontal axis indicates log2 fold change, while vertical axis indicates adjusted p-value of RA vs HD differences. Point sizes indicate average gene expression level (normalized counts). Significantly up- or downregulated genes (adjusted p-value < 0.05, absolute log2FC > 1) are color-coded red or blue respectively. Significant genes with absolute log2FC > 3 are labeled to aid interpretation. Upregulated genes include inflammatory chemokines, immediate early genes, and extracellular matrix–related genes, while downregulated genes include ST8SIA6 and metabolic regulators. **(E)** Gene Set Enrichment Analysis (GSEA) performed using the t statistic of RA vs HD untreated M0 monocytes differential gene expression analysis. Point colors indicate normalized enrichment score (NES) whereby positive and negative scores indicate up- and downregulation in RA respectively, while point sizes indicate number of leading edge genes. The vertical dashed line indicates the adjusted p-value = 0.05 threshold. Top 15 gene sets with smallest adjusted p-value are shown. Top gene sets are enriched with inflammatory pathways (HALLMARK_INFLAMMATORY_RESPONSE, TNFA_SIGNALING_VIA_NFKB, Signaling by Interleukins) and the KEAP1-NFE2L2 oxidative stress pathway, and downregulation of Translation, E2F targets, and Mitochondrial matrix gene sets.

To assess the biological relevance of these modules, we performed ORA on genes assigned to each module. Among the most enriched gene sets per module, defined by maximal gene overlap (**Fig. 2A, right**), several expected patterns emerged.

*HALLMARK_INTERFERON_GAMMA_RESPONSE* was the top enriched gene set for the ‘green’ module, which was upregulated in M(IFNγ+LPS) and downregulated in M(IL-4), and for the ‘pink’ module, which was upregulated following acute LPS stimulation. In addition, the ‘blue’ module, which was upregulated in M(IFNγ+LPS), and the ‘brown’ module, which was upregulated in M(IL-4) and downregulated by LPS, were enriched for *GO:0005759 mitochondrial matrix*, consistent with differential mitochondrial engagement across macrophage states.

Among the modules with RA-associated correlations, the ‘black’, ‘salmon’, and ‘darkred’ showed modest correlations with disease status (-0.45, -0.42, and 0.31, respectively; **Fig. 2A, middle**). The ‘black’ module was enriched for *GO:0002250 adaptive immune response*, albeit these genes were downregulated, consistent with monocytes being primarily a component of innate rather than adaptive immunity. The ‘salmon’ and ‘darkred’ modules showed enrichment for *R-HSA-72766 Translation* and *MIR1233_3P*, respectively. Examination of overlapping genes between the ‘darkred’ module, which was positively correlated with both RA and M(IFNγ+LPS) differentiation, and the *MIR1233_3P* gene set revealed that this overlap consisted almost exclusively of protocadherin family members of the *PCDHGA* and *PCDHGB* clusters (**Fig. 2B**). This suggests that protocadherin genes are upregulated in two inflammation-associated contexts. Notably, the ‘darkred’ module itself was composed almost entirely of clustered protocadherin genes, many of which were annotated as predicted targets in multiple miRNA target gene sets. Because the module did not include other genes expected within these target sets, the apparent enrichment likely reflects the recurrent inclusion of highly homologous protocadherin genes in target prediction databases rather than specific miRNA-mediated regulation.

In contrast, the ‘salmon’ module, which was negatively correlated with RA but positively correlated with M(IFNγ+LPS), was dominated by ribosomal proteins and translation initiation or elongation factors (**Fig. 2C**). This pattern points to divergence in translational regulation between RA monocytes and classically activated macrophages, indicating that RA monocytes do not simply represent an M(IFNγ+LPS)-like inflammatory state. Rather, classically activated macrophages only partially recapitulate the transcriptional changes associated with RA.

Differential expression analysis of untreated samples within each differentiation state revealed a focused RA-associated transcriptional program in M0 monocytes (**Fig. 2D**). Upregulated genes included inflammatory chemokines (e.g., *CCL3*, *CXCL8*), immediate early response genes (*FOS*, *EGR1*), and extracellular matrix-related genes (e.g., *TNC*, *ADAMTS2*). Additional genes with strong statistical significance included *EME2*, *TBC1D3K*, and *ZNF316*. RA monocytes also showed increased expression of genes linked to neuronal or endocrine pathways (e.g., *GAL*, *TRIM9*) as well as hemoglobin subunit *HBG1*. Downregulated genes included regulators of endolysosomal function and vesicle trafficking (e.g., *ELAPOR2*, *AGAP1*), metabolic enzymes (*ALDH7A1*), and glycosylation-related genes such as *ST8SIA6*.

M(IFNγ+LPS) macrophages showed more extensive RA-associated transcriptional changes than M0 monocytes (**Fig. S2D**), including upregulation of inflammatory chemokines (e.g., *CCL17*, *CXCL1*), cytokines (*IL24*), protease inhibitors (*SERPINB* family), and lectins (*MRC1*, *CLEC5A*), as well as genes linked to inflammation and tissue remodeling such as *SPP1*.

Additional changes included genes involved in polyamine metabolism (*AOC1*) and glycoproteins (*MUCL1*). Downregulated genes comprised metabolic enzymes (e.g., *SHMT2*, *ASS1*) and immunoregulatory factors (*ORM1/2*). In contrast, M(IL-4) macrophages exhibited minimal RA-associated changes (**Fig. S2E**), with only a small set of differentially expressed genes, including *SLC16A9*, *UCP3*, and *NR6A1*, and limited overlap with other states (**Fig. S2F**), indicating that RA-associated transcriptional alterations are context-dependent.

GSEA in M0 monocytes identified upregulation of inflammatory and stress-related pathways, including TNF/NF-κB and interleukin signaling, alongside oxidative stress responses (**Fig. 2E**). In parallel, pathways related to cell cycle, translation, and mitochondrial function were downregulated, consistent with the WGCNA-derived translation module and indicating constrained biosynthetic capacity.

These findings were validated in CD14+ monocytes from an independent cohort from the RA-MAP project (25) which showed consistent upregulation of inflammatory pathways and downregulation of translation-related programs despite differences in sample handling (**Fig. S2H**), supporting the robustness of the observed transcriptional signature.

In summary, while differentiation state dominated transcriptomic variation, RA was associated with a focused transcriptional program characterized by inflammatory activation, reduced translation and mitochondrial pathways, and a distinct protocadherin-associated signature.

### Proteome analysis identifies rheumatoid arthritis-associated perturbations of the secretory pathway

For the proteomics dataset, we applied an analysis workflow analogous to that used for transcriptomics. As in the transcriptomic data, differentiation state (M0, M(IFNγ+LPS), and M(IL-4)) was the primary driver of clustering (**Fig. S3A**) and accounted for the largest fraction of overall variance, with donor identity representing the second strongest contributor (**Fig. S3B**).

When analyzing each differentiation state separately, donor-specific effects dominated, whereas acute LPS treatment contributed to variance within the M0 and M(IL-4) subsets (**Fig. S3C**). This is consistent with prior LPS exposure during the generation of M(IFNγ+LPS) macrophages, which likely reduces responsiveness to subsequent acute stimulation.

WGCNA identified 16 modules in the proteomic dataset. Most modules correlated strongly with M(IFNγ+LPS) and/or M(IL-4) differentiation, whereas two modules (‘midnightblue’ and ‘magenta’) were associated with LPS treatment (r = 0.49 and 0.63, respectively). No module showed a strong correlation with RA disease status, with the highest correlation observed for the ‘greenyellow’ module (r = -0.30) (**Fig. 3A, middle**).

**Figure 3.**
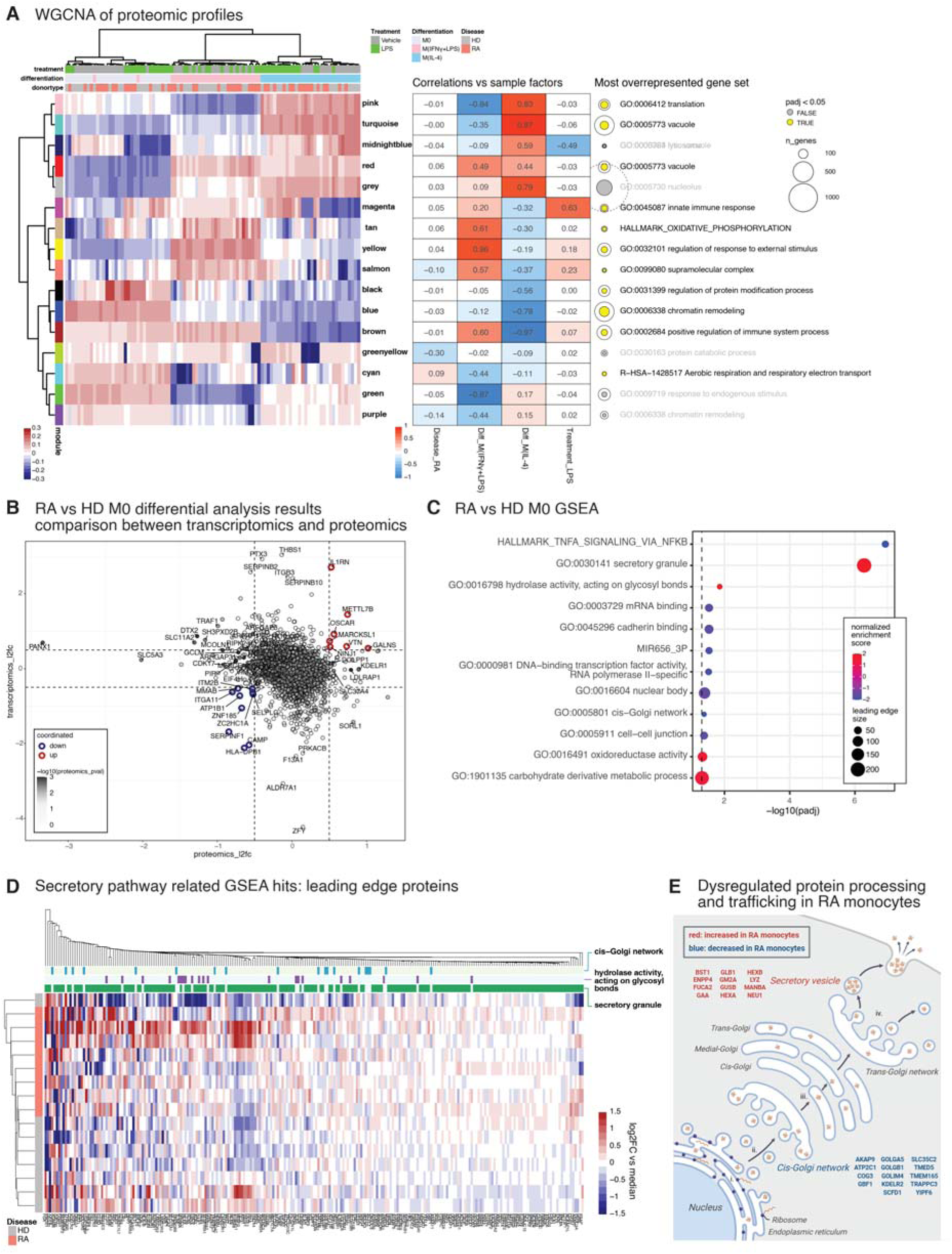
Proteome analysis identifies rheumatoid arthritis-associated perturbations of the secretory pathway. **(A)** Weighted correlation network analysis (WGCNA) of proteomics data identifying 16 protein modules. Left: clustered heatmap showing module eigengene values in individual samples, as well as clustering patterns of samples according to eigengenes. Center: heatmap of correlations between module eigengenes and sample traits including disease status (RA, HD), differentiation state (M0, M(IFNγ+LPS), M(IL-4), and acute LPS treatment. Right: top overrepresented gene set, defined as having the largest number of overlap genes, for each module. The double concentric circles next to gene set names represent number of genes in the module (size of outer circle) and number of overlap genes (size of inner circle), respectively. The inner circle is color coded by statistical significance of the top overrepresented gene set (yellow if adjusted p-value < 0.05, grey otherwise). Differentiation state dominates module associations, whereas no module shows a strong correlation with RA. **(B)** Scatter plot comparing RA vs HD log2 fold-changes from transcriptomic and proteomic differential analyses in untreated M0 monocytes. The transcriptomics differential analysis compared 6 RA vs 6 HD, while the proteomics analysis compared 8 RA vs 8 HD samples; all samples were biologically independent. The vertical and horizontal axes represent transcriptomics and proteomics log2FCs respectively. Data points are additionally shaded according to proteomics RA vs HD p-value. Genes up- or downregulated consistently in both transcriptomics and proteomics (above absolute log2FC threshold of 0.5) are outlined red or blue, respectively. Specific genes are labeled to aid interpretation. Consistently upregulated genes include *IL1RN*, *OSCAR*, *VTN*, *NINJ1*; downregulated genes include *SELPLG*, *ITGA11*, *ATP1B1*, *HLA-DPB1*, *MMAB*. **(C)** Gene Set Enrichment Analysis (GSEA) performed using the t statistic of RA vs HD untreated M0 monocytes differential protein abundance analysis. Point colors indicate normalized enrichment score (NES) whereby positive and negative scores indicate up- and downregulation in RA respectively, while point sizes indicate number of leading edge genes. The vertical dashed line indicates the adjusted p-value = 0.05 threshold. 12 gene sets with adjusted p-value < 0.05 are shown. Upregulated gene sets include *GO:0030141 secretory granule* and *GO:0016798 hydrolase activity, acting on glycosyl bonds*; downregulated gene sets include *GO:0005801 cis-Golgi network* and *HALLMARK_TNFA_SIGNALING_VIA_NFKB*. **(D)** Clustered heatmap of leading-edge proteins from secretory pathway–related gene sets (*GO:0030141 secretory granule*, *GO:0016798 hydrolase activity, acting on glycosyl bonds* and *GO:0005801 cis-Golgi network*) demonstrating coordinated RA-associated abundance shifts. Protein abundances are expressed as log2 fold-change over median abundance of the respective protein within the dataset. **(E)** Schematic illustration summarizing protein abundance perturbations within the secretory pathway in RA monocytes: depletion of cis-Golgi–associated proteins and accumulation of distal secretory vesicle proteins.

We next examined RA - HD differences at the level of individual proteins. Due to substantial within-group variability and missing values, differential analysis of untreated monocytes did not identify proteins meeting an adjusted p-value threshold of 0.05. We therefore focused on genes showing concordant changes at both transcriptomic and proteomic levels. Using an absolute log2 fold-change threshold of 0.5, we identified a set of genes with consistent RA-associated differences in M0 monocytes (**Fig. 3B**).

Upregulated proteins included IL1RN, encoding the interleukin-1 receptor antagonist; *METTL7B*, an alkyl thiol methyltransferase implicated in RA; *OSCAR*, a co-stimulatory receptor required for osteoclast differentiation; *VTN* (vitronectin), a mediator of cell adhesion and spreading; GALNS, a lysosomal enzyme involved in glycosaminoglycan degradation; *NINJ1*, a mediator of plasma membrane rupture during lytic cell death; and *MARCKSL1*, which regulates actin cytoskeleton dynamics. Downregulated proteins included *SELPLG* (PSGL-1), *ITGA11*, and *ZNF185*, all involved in cell adhesion; *ITM2B*, associated with intracellular trafficking; *MMAB*, a mitochondrial enzyme involved in vitamin B12 metabolism; *ATP1B1*, a subunit of the Na⁺/K⁺ ATPase; *CAMP*, encoding the antimicrobial peptide cathelicidin; *HLA-DPB1*, a component of MHC class II; and *ZC2HC1A*.

As observed in the transcriptomic dataset, M(IFNγ+LPS) and M(IL-4) macrophages exhibited distinct RA-associated changes relative to M0 monocytes (**Fig. S3D, E**). MRC1 was the most strongly upregulated gene in M(IFNγ+LPS) macrophages from RA patients at both transcript and protein levels. Additional upregulated proteins in these cells included *THBS1*, *A2M*, *NRP1*, *PZP*, and *CD109*, whereas *CD163* was downregulated. In M(IL-4) macrophages, RA-associated increases included *SLC2A1*, *PFKM*, *THBS1*, *FSCN1*, *PROS1*, *HBA2*, *HBB*, and SERPINF1, while *APOL3*, *NMT2*, and *MMAB* were decreased (**Fig. S3E**). Notably, *MMAB* was consistently downregulated across M0 and M(IL-4) cells, whereas *THBS1* was upregulated in both M(IFNγ+LPS) and M(IL-4) macrophages.

Given the modest differences at the level of individual proteins, we next assessed pathway-level changes using GSEA based on RA-HD differential analysis in untreated monocytes (**Fig. 3C**). Upregulated gene sets included *GO:0030141 secretory granule* and *GO:0016798 hydrolase activity, acting on glycosyl bonds*. These gene sets represent proteins localized to distal compartments of the secretory pathway and enzymes with glycosidase or glycosyltransferase activity, respectively. Shared genes included *MANBA*, *FUCA2*, *GAA*, *GLB1*, *GUSB*, *HEXA*, and *LYZ* (Fig. S3F). In contrast, *GO:0005801 cis-Golgi network*, representing proximal components of the secretory pathway, was among the downregulated gene sets.

A clustered heatmap of proteins within these gene sets showed clear separation between RA and HD samples despite modest changes at the level of individual proteins (**Fig. 3D**). Together, these findings indicate a shift in the secretory pathway, characterized by reduced abundance of proteins associated with early compartments and increased abundance of proteins associated with distal secretory vesicles (**Fig. 3E**).

Unexpectedly, we observed downregulation of the *HALLMARK_TNFA_SIGNALING_VIA_NFKB pathway* at the protein level, including canonical components such as *NFKB1*, *NFKB2*, *JUNB*, *PTPRE*, *NAMPT*, and *SQSTM1* (**Fig. S3F**), despite transcriptional upregulation of this pathway (**Fig. 2E**). Although fold changes were modest, hierarchical clustering of these proteins was sufficient to distinguish RA from HD samples, with a consistent trend toward reduced protein abundance in RA (**Fig. S3G**). Additional downregulated gene sets included *GO:0000981 DNA-binding transcription factor activity, RNA polymerase II-specific*; *GO:0003729 mRNA binding*; *GO:0045296 cadherin binding*; *GO:0016604 nuclear body*; and *GO:0005911 cell–cell junction*.

In summary, proteomic analysis revealed limited RA-associated differences at the level of individual proteins but identified consistent pathway-level alterations. These included perturbations of the secretory pathway and a discordance between transcriptional and protein-level regulation of NFκB signaling.

### Multi-omics integration identifies rheumatoid arthritis-associated reductions in glycosylation-related metabolic fluxes

Beyond single-omics analyses, integrative multi-omics approaches can uncover relationships between gene expression, protein abundance, and metabolic phenotype. We therefore performed integrated analyses across transcriptomic, proteomic, and metabolomic datasets.

We first integrated transcriptomics and metabolomics at the pathway level using IMPaLA, which combines enrichment scores from independent differential analyses. Pathways with the highest joint significance included translation, metabolism of amino acids and derivatives, selenoamino acid metabolism, metabolism of RNA, and aminoacyl-tRNA biosynthesis (**Fig. 4A**). Integration of proteomics and metabolomics identified additional pathways, including diabetic cardiomyopathy, post-translational protein modification, and metabolism of carbohydrates and lipids (**Fig. 4B**).

**Figure 4.**
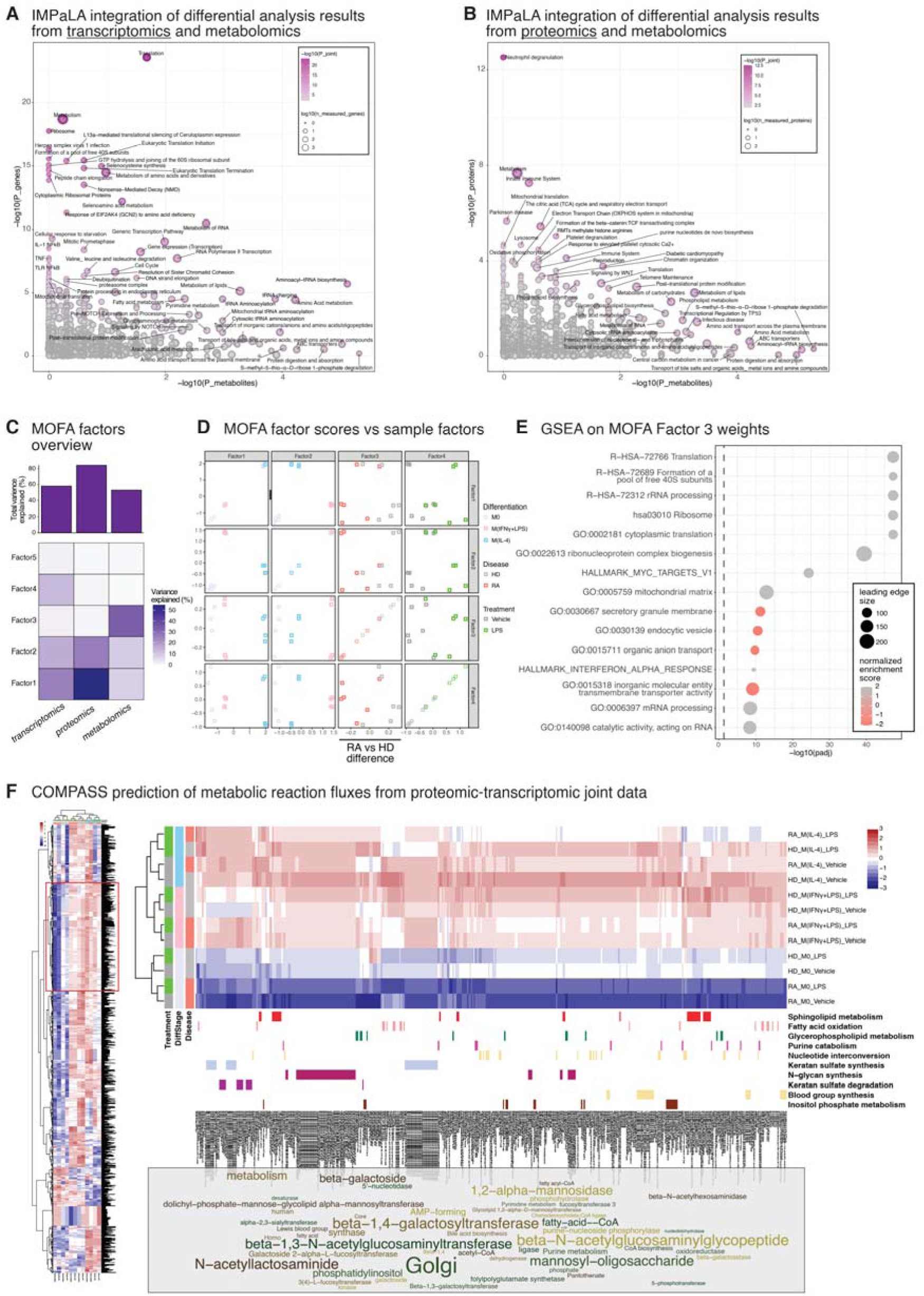
Multi-omics integration identifies rheumatoid arthritis-associated reductions in glycosylation-related metabolic fluxes. **(A)** Integrated Molecular Pathway Level Analysis (IMPaLA) identifying metabolic pathways jointly enriched across transcriptomics and metabolomics datasets. The scatter plot shows p-values of enrichment based on individual datasets – transcriptomics (vertical axis) or metabolomics (horizontal axis) – as well as p-value of joint enrichment indicated by data point color. The data point sizes indicate number of measured genes involved in each metabolic pathway. Jointly enriched pathways include Translation, Amino acid metabolism, Metabolism of RNA, and Aminoacyl-tRNA biosynthesis. **(B)** IMPaLA identifying metabolic pathways jointly enriched across proteomics and metabolomics datasets. The scatter plot shows p-values of enrichment based on individual datasets – proteomics (vertical axis) or metabolomics (horizontal axis) – as well as p-value of joint enrichment indicated by data point color. The data point sizes indicate number of measured genes involved in each metabolic pathway. Jointly enriched pathways include Diabetic cardiomyopathy, Post-translational protein modification, Carbohydrate metabolism, and Lipid metabolism. **(C)** Overview of Multi-Omics Factor Analysis (MOFA) model using pooled pseudosamples across transcriptomics, proteomics and metabolomics datasets. Upper panel: total variance explained per omics layer. Lower panel: variance explained per omics layer, per MOFA factor. **(D)** Scatterplots visualizing the association of MOFA factors with sample traits. Data points in each scatterplot represent pseudosamples and are color coded according to either differentiation (M0 / M(IFNγ+LPS) / M(IL-4), disease status (HD / RA) or LPS treatment (Vehicle/ LPS). Factor 1 corresponds to M(IL-4), Factor 2 to M(IFNγ+LPS), Factor 3 to disease status (HD positive, RA negative), and Factor 4 to acute LPS treatment. **(E)** Gene Set Enrichment Analysis (GSEA) performed on MOFA Factor 3, which corresponds to separation of HD vs RA pseudosamples. For each gene, the corresponding transcript and protein MOFA Factor 3 weights are summed and used as the ranking statistic. Hence, genes with both transcript and protein weights in the same direction (positive or negative) will strengthen the gene’s contribution to the GSEA, while opposing directions between transcript and protein weights will decrease the gene’s overall weight in the analysis. Point colors indicate normalized enrichment score (NES) whereby positive and negative scores indicate up- and downregulation in RA respectively, while point sizes indicate number of leading edge genes. The vertical dashed line indicates the adjusted p-value = 0.05 threshold. Top 15 gene sets with smallest adjusted p-value are shown. Positive enrichment (higher in HD, decreased in RA) includes Translation, Ribosome biogenesis, Mitochondrial matrix, and MYC targets; negative enrichment (increased in RA) includes Secretory granule membrane and Endocytic vesicle pathways. **(F)** Reaction scores predicted by Compass metabolic modeling on combined transcriptomics-proteomics merged pseudosamples. The overall heatmap containing all reactions is shown as a thumbnail on the left. The main center panel shows a zoomed-in section of the heatmap with a cluster of reactions predicted to have decreased flux in RA M0 monocytes. Metabolic subsystems (*Sphingolipid metabolism*, *Fatty acid oxidation*, *Glycerophospholipid metabolism*, …) corresponding to reactions in this cluster are indicated. The bottom part shows a word cloud illustration of the terms appearing most frequently in the reaction names of this cluster, highlighting that Golgi and glycosylation related reactions are highly enriched in this cluster.

We next performed integrative analysis across omics layers using Multi-Omics Factor Analysis (MOFA) (26). As the datasets were generated from independent donors, we applied a pseudosample approach in which samples sharing the same combination of disease status (RA vs HD), differentiation state (M0, M(IFNγ+LPS), M(IL-4)), and treatment (vehicle vs LPS) were averaged within each omics layer and treated as matched across datasets (**Fig. S4A**). A MOFA model with five factors was constructed.

The model explained 57.7%, 83.8%, and 52.6% of the variance in transcriptomic, proteomic, and metabolomic data, respectively (**Fig. 4C, upper panel**). Factors 1 and 2 were primarily driven by proteomic variation and corresponded to M(IL-4) and M(IFNγ+LPS) differentiation states, respectively (**Fig. 4C, D**). Factor 3 separated disease status, with positive scores associated with HD and negative scores with RA. Factor 4 reflected acute LPS treatment, whereas factor 5 explained minimal variance and was not further considered.

To characterize disease-associated programs, we performed GSEA on Factor 3 weights using the combined transcriptomic and proteomic contributions for each gene. Gene sets with positive enrichment scores (decreased in RA) largely recapitulated pathways identified in single-omics analyses, including cytoplasmic translation, ribosome biogenesis, rRNA processing, and mitochondrial matrix (**Fig. 4E; Fig. S4B**). In addition, *HALLMARK_MYC_TARGETS_V1* was significantly enriched, driven by genes involved in nucleotide metabolism and cell cycle regulation.

Gene sets with negative enrichment scores (increased in RA) included secretory granule membrane and endocytic vesicle components, consistent with proteomic evidence for altered intracellular trafficking. Additional enriched pathways included organic anion transport and transmembrane transporter activity, suggesting reduced expression of transporters across transcript and protein levels.

To further investigate metabolic consequences, we integrated transcriptomic and proteomic data using the Compass algorithm (27) to predict metabolic reaction feasibility. When protein abundance was available, it was used to estimate enzyme activity; otherwise, transcript levels were used as proxies (see Methods). Using pseudosamples, Compass identified a cluster of reactions consistently decreased in both vehicle- and LPS-treated RA monocytes relative to HD controls (**Fig. 4F**).

Annotation of these reactions revealed a strong enrichment for glycosylation-related processes, including reactions associated with Golgi function, glycosyltransferases (e.g. beta-1,4-galactosyltransferase and beta-1,3-N-acetylglucosaminyltransferase), and glycan synthesis pathways. These reactions mapped to metabolic subsystems such as *N-glycan biosynthesis*, *keratan sulfate metabolism*, *sphingolipid metabolism*, *glycerophospholipid metabolism*, and *blood group antigen synthesis*. Additional affected subsystems included *fatty acid oxidation*, *inositol phosphate metabolism*, *nucleotide interconversion*, and *purine catabolism*.

To exclude potential artifacts introduced by the pseudosample approach, we repeated Compass analysis using individual transcriptomic and proteomic samples followed by RA–HD differential analysis of reaction scores (**Fig. S4C**). This analysis confirmed reduced predicted feasibility of glycosylation-related reactions in RA, supporting the robustness of the findings.

Together, these integrative analyses indicate that RA monocytes exhibit coordinated alterations across multiple layers of regulation, including reduced translation-related programs, perturbations in intracellular trafficking, and decreased glycosylation-associated metabolic activity.

## DISCUSSION

In this study, we performed an integrated multi-omics analysis of primary human monocytes from RA patients and healthy controls under controlled differentiation and inflammatory conditions. A central finding is the striking divergence in how disease status manifests across molecular layers. Whereas transcriptomic and proteomic variation was primarily driven by differentiation state and acute LPS stimulation, variance in the metabolomic data was most strongly associated with RA disease status. These findings indicate that circulating RA monocytes have a distinct metabolic phenotype that is largely independent of experimental stimulation. This is consistent with a model in which circulating RA monocytes exist in a metabolically pre-programmed state that may shape their functional responses and differentiation trajectories.

Our study recovered many expected features of RA monocyte and macrophage biology, including inflammatory and interferon-associated transcriptional programs, differentiation-dependent mitochondrial remodeling, and well-established immunometabolic signatures such as succinate accumulation in inflammatory contexts. The recovery of these canonical signals across transcriptomics, proteomics, and metabolomics supports the validity of the experimental design and analytical framework. In this context, novel observations – particularly those identified at the level of modules or pathways rather than individual features – are more likely to reflect bona genuine disease-associated biology rather than technical noise.

Our findings suggest that RA monocytes are not only characterized by increased inflammatory signaling but also by broader metabolic reprogramming. From metabolomics data, we found that undifferentiated monocytes as well as differentiated M(IFNγ+LPS) and M(IL-4) cells from RA patients consistently exhibited broad alterations across amino acid metabolism, central carbon metabolism, lipid metabolism, nucleotide metabolism, and redox-related pathways. The coordinated depletion of nucleotides, nucleotide sugars, and glycerophospholipid intermediates in RA monocytes was particularly striking. These metabolites are essential substrates for biosynthesis, membrane remodeling, and protein glycosylation. Their reduction suggests a metabolic bottleneck that could have far-reaching consequences for protein processing and secretion. Indeed, impacts on protein processing and secretion may be indirectly linked to another observed metabolic aspect: broad accumulation of amino acids in RA samples, which led to strong enrichment of the *hsa00970 Aminoacyl−tRNA biosynthesis* KEGG pathway in both WGCNA-ORA and differential analysis-GSEA. Specifically, disrupted protein processing and secretion would be expected to feed back into regulatory mechanisms controlling protein synthesis rate, impacting protein translation (28–30). This may potentially explain the prominent signature of transcriptomic downregulation of protein translation related gene sets.

WGCNA also identified two metabolite modules corresponding to the proximal or upstream, and distal or downstream parts of the kynurenine pathway. Both modules increased with M(IFNγ+LPS) differentiation and acute LPS challenge, but showed differential correlation with RA disease status: while the module containing proximal intermediates (kynurenine, 3-hydroxyanthranilate) was increased in RA, the module containing distal metabolites (picolinate, quinolinate) was decreased (**Fig. 1A**). A recent mechanistic study demonstrated that inflammatory activation induces upstream kynurenine pathway metabolites while limiting distal conversion to NAD⁺, thereby promoting a pro-inflammatory macrophage phenotype (31). Our data in RA monocytes closely mirror this metabolic state, with accumulation of IFN-associated kynurenine metabolites and reduced NAD⁺-related metabolites, suggesting that circulating RA monocytes are pre-activated in the bloodstream and display a metabolic phenotype characteristic of inflammatory activation.

Notably, metabolic responses to differentiation and LPS-stimulation were less prominent in RA monocytes compared to HDs. In HD monocytes, differentiation and LPS-stimulation produced the expected changes in metabolites such as increased itaconate and kynurenine pathway intermediates, whereas these responses appeared blunted in RA samples (**Fig. S1A**). These findings suggest that RA monocytes exist in a metabolically constrained or preconditioned state that limits their ability to further remodel metabolism in response to differentiation or inflammatory cues.

We considered the possibility that RA monocytes exhibited depressed nucleotide and nucleotide sugar levels due to nucleotide-targeting drugs such as methotrexate or leflunomide, which are common treatments for RA. However, we did not observe any association between patient methotrexate or leflunomide treatment and the metabolic signature of RA monocytes in both discovery and validation cohorts (**Fig. S1G, Fig. S1I**). The consistent observation of depressed nucleotide metabolite levels in RA patient monocytes across two cohorts, despite differences in sample processing between cohorts (24 hours resting post-isolation for the discovery cohort and immediate harvesting for validation cohort) supports the notion that this is a disease-intrinsic metabolic difference already present in circulating monocytes, rather than a treatment effect or secondary consequence of transitioning to ex vitro culture.

At the transcriptomic level, RA-associated effects were modest in magnitude and restricted to a subset of genes. Nonetheless, WGCNA and GSEA revealed coherent disease-associated modules involving protein translation, mitochondrial matrix components, and protocadherin family genes (**Fig. 2A, 2E**). Inspection of the GSEA leading edge genes for the *GO:0005759 mitochondrial matrix* gene set (**Fig. S2G**) revealed predominantly metabolic genes among the top genes (i.e. showing strongest, most consistent downregulation in RA), including those involved in TCA cycle flux and anaplerosis (*IDH2, GOT2, SDHAF3/4*), branched-chain amino acid oxidation (*BCKDHB, DBT, BCAT2*), fatty acid oxidation (*HADH, ACACB, ACSF2*), and aldehyde metabolism (*ALDH6A1, ALDH7A1*), and heme biosynthesis (*FECH*). Other top genes included components of the mitochondrial translation apparatus (*MRPS27, MRPS18B, MRPL30, DARS2, IARS2, NARS2*), mitochondrial sirtuins (*SIRT4, SIRT5*) and redox-associated enzymes (*IDH2, NUDT1, GSTK1*), implicating disruption of NAD⁺-dependent regulatory circuits and redox buffering systems. These findings align with our metabolomics data showing disease-associated decreases in NAD(H) and NADP(H) levels, supporting a model whereby reduced NAD(H) availability constrains mitochondrial metabolic fluxes, limits antioxidant capacity, and dampens sirtuin-mediated post-translational control in RA monocytes.

The transcriptional downregulation of ribosomal and translation-related gene sets in RA monocytes despite slight upregulation under M(IFNγ+LPS) polarization (**Fig. 2A**) suggests that RA monocytes appear to occupy a distinct state from canonical inflammatory activation, in which protein translation programs are selectively dampened. On the other hand, protocadherin genes were upregulated in both RA and M(IFNγ+LPS) (**Fig. 2B**), potentially pointing to an inflammation-related signature. Clustered protocadherins (PCDHA/B/C) are a large subfamily of cadherin-superfamily cell adhesion proteins whose expression is tightly regulated in neurons, and are crucial for neuronal self-recognition during development (32–34). Although roles in classical immune/inflammatory pathways are not established, protocadherin family members have been implicated in cell migration and adhesion in myeloid cells and other contexts (35), suggesting that regulated adhesion or signaling could modulate macrophage functions.

Several individual RA-associated transcript changes implicate perturbations in protein trafficking and post-translational modification. Differential expression of genes such as *TBC1D3K*, *AGAP1*, *NEU4*, and *ST8SIA6* (**Fig. 2D**) suggests altered vesicle trafficking, endocytosis, lysosomal processing, and sialylation capacity – all supporting a general theme of dysregulated secretory pathway organization and/or function. Although these genes have not been widely studied in RA monocytes, they converge functionally on intracellular transport and glycosylation pathways.

Proteomic analyses further supported the notion that RA monocytes exhibit perturbations in intracellular trafficking and secretion. While no individual proteins reached statistical significance after multiple-testing correction, coordinated changes were evident at the pathway level. Gene set enrichment analysis revealed enrichment of secretory granule and glycosyl hydrolase–related proteins alongside depletion of cis-Golgi network components (**Fig. 3C**), altogether pointing to altered secretory pathway organization. Considering the central importance of the secretory pathway compartments such as ER and Golgi for cellular glycosylation, it is remarkable that constraint-based metabolic modeling further predicted widespread reductions in glycosylation-related metabolic fluxes in RA monocytes (**Fig. 4F**, **Fig. S4C**). Together with the depleted nucleotide sugar levels observed in metabolomics data, these findings support a model of metabolic–secretory decoupling, in which reduced metabolic capacity constrains protein processing and glycosylation despite ongoing inflammatory activation.

The protein glycosylation-secretion axis is increasingly recognized as a critical regulator of immune function. Protein secretion and vesicular transport pathways govern the release of cytokines, proteases, and extracellular matrix–modifying enzymes, while glycosylation modulates immune receptor signaling and protein stability (36–39). Altered glycosylation of IgG – particularly reduced galactosylation and sialylation – is a well-established hallmark of RA and contributes to enhanced Fc receptor–mediated inflammation (40–42). However, altered glycosylation on immune cells themselves are far less well-studied, particularly for cells of the myeloid lineage or in the context of autoimmune disease. Our findings suggest that analogous alterations in glycosylation capacity may occur within monocytes and macrophages themselves, potentially influencing receptor signaling, cytokine secretion, and cell-cell interactions. Perturbations in vesicular trafficking and secretion pathways may further amplify these effects by altering the processing and release of inflammatory mediators, chemokines, and matrix-modifying enzymes.

Another interesting finding from the proteomics GSEA was the observed downregulation of NF-κB pathway components, which stands in stark contrast to the clear transcriptomic upregulation of the very same pathway. This finding underscores the often-overlooked reality that steady-state protein abundance does not necessarily mirror transcriptional activation, and expression changes at the protein level may even oppose transcriptional differences due to post-transcriptional regulation as had been previously reported (43, 44). In particular, reviews of NF-κB pathway regulation note that extensive post-transcriptional control (mRNA stability, translation control) exists within this pathway, providing mechanistic basis for decoupled transcriptional and proteomic responses (45).

Several RA-associated proteomic changes point to functional consequences for tissue inflammation and remodeling. For example, upregulation of *THBS1* at both transcript and protein levels in differentiated RA macrophages is notable given its role as a multifunctional secreted glycoprotein that modulates cell adhesion, extracellular matrix interactions, and immune receptor signaling (46, 47). Similarly, differential abundance of adhesion molecules, scavenger receptors, and cytoskeletal regulators suggests altered migratory and tissue-interactive properties of RA monocytes and macrophages. The downregulation of *CD163* in RA-derived M1 macrophages is consistent with prior evidence that CD163 has protective roles in arthritis models, and its depletion may exacerbate inflammatory pathology (48, 49). Here, we acknowledge that throughout our analyses we have largely focused on RA vs HD differences in M0 and have not fully explored RA vs HD differences in M(IFNγ+LPS) and M(IL-4) cells. Future detailed analyses of our dataset, focused on these additional comparisons, are expected to yield further insight into the influence of RA disease status on inflammatory activation of monocyte-derived macrophages.

Several limitations of this study merit consideration. Due to practical constraints, transcriptomic, proteomic, and metabolomic data were generated from independent sample sets rather than fully matched samples, necessitating pseudosample pooling for certain integrative analyses. While multiple orthogonal approaches yielded convergent results, inspiring confidence in the validity of our findings, future studies with fully matched multi-omics measurements at the single-donor level will doubtless refine and deepen the insights obtained in this study. In addition, the use of circulating monocytes, as well as simulation of inflammatory activation by in vitro differentiation protocols and LPS challenge cannot be expected to fully recapitulate the complexity of inflammatory responses within the synovial microenvironment. Nonetheless, we found value in our controlled experimental design that enabled clear separation of disease-, differentiation-, and stimulation-associated effects, which is difficult to achieve in tissue-based studies.

In summary, integrated analysis of our multi-omics dataset reveals that RA monocytes are characterized by systemic metabolic reprogramming that is also partially reflected at the transcriptomic and proteomic levels. This metabolic–secretory decoupling links reduced nucleotide, redox, and glycosylation resources to altered protein processing and intracellular trafficking. Together, this study highlights metabolic–secretory decoupling as a promising axis for future mechanistic investigation and therapeutic intervention in RA, while also providing a resource for hypothesis generation on monocyte immune dysfunction in RA.

## METHODS

### Study participants and ethics

Ethical approval for this study was obtained from the Ethikkommission Nordwest- und Zentralschweiz (EKNZ; EKNZ 2019-01693) and the Kantonale Ethikkomisison Zürich (KEK; KEK-ZH-NR. 515). The study was conducted in accordance with the Declaration of Helsinki, and written informed consent was obtained from all participants prior to enrolment. Patients with seropositive rheumatoid arthritis (RA) in the discovery cohort (metabolomics, transcriptomics and proteomics) were recruited at the Department of Rheumatology, University Hospital Basel, Switzerland and an independent validation cohort of RA patients was recruited at the Department of Rheumatology, University Hospital Zurich for metabolomics (**Fig. S1G**). All patients fulfilled the 2010 ACR/EULAR Rheumatoid Arthritis Classification Criteria and were required to be seropositive for anti-citrullinated protein antibodies (ACPAs). Patient enrolment was restricted to ACPA-positive individuals to obtain a more homogeneous RA patient population. Peripheral blood samples were obtained during routine outpatient visits. Clinical disease activity was assessed during the same visit using DAS-28-CRP scoring. Exclusion criteria included the presence of another autoimmune disease, active infection, malignancy, or systemic corticosteroid treatment within the preceding 14 days. Age- and sex-matched healthy donors were recruited through the Blutspende SRK Nordwestschweiz blood donation service (discovery cohort) and the Blutspende SRK Zürich, Spendezentrum Schlieren.

### Isolation of primary human CD14+ monocytes (M0) from buffy coat preparations

Isolation of peripheral blood mononuclear cells (PBMCs) was done by standard density-gradient centrifugation (Lymphoprep, Fresenus Kabi, Norway). Positive selection of monocytes (M0) using CD14 Microbeads (Miltenyi Biotec, Germany) was performed according to manufacturer’s instructions. Isolated monocytes were cultured in R10 media (complete medium (RPMI 1640 (Gibco) with 10% heat-inactivated fetal bovine serum (FBS) and 1% penicillin- streptomycin supplemented with 2% GlutaMAX (all from GIBCO, ThermoFisher Scientific). Cell density and viability were determined by Trypan Blue staining. Trypan Blue 0.4% (Nano EnTek) was mixed 1:1 with 10 μl cell suspension and pipetted on a counting slide (Nano EnTek). Counting was performed by an EVE^TM^ Automated Cell Counter (NanoEnTek).

### Cell culture & metabolite extraction for metabolomics

Differentiation of CD14+ monocytes into classically and alternatively activated macrophages was achieved by supplementing the culture with 20 ng/mL LPS/IFNγ or 20 ng/mL IL-4, respectively, on a three-times-per-week basis for 8 to 11 days. For macrophage activation, a 24-hour stimulation with 100 ng/mL LPS was used. Following a 24-hour incubation, the cell culture media was aspirated, and the cells were washed twice with ice-cold PBS. After the final wash, PBS was completely removed, and metabolites were extracted by the addition of a solution consisting of 50% LC-MS grade Methanol (Fisher Scientific, 10284580), 30% LC-MS grade Acetonitrile (Fisher Scientific, 10001334), and 20% ultrapure water. This extraction process was performed on dry ice, and the plates were subsequently stored at -80°C overnight. The following day, the cell extraction suspension was collected into pre-cooled Eppendorf tubes and placed in a thermoshaker at 4°C, operating at maximum speed for 15 minutes. Samples were then centrifuged for 20 minutes at 4°C at 15,000 rpm. The upper 80% of the supernatant was carefully collected and transferred into autosampler vials for analysis.

### Untargeted LC-MS/MS metabolomics

Metabolites were separated chromatographically using a Millipore SeQuant ZIC-pHILIC analytical column (5 µm, 2.1L×L150Lmm) complemented by a 2.1L×L20Lmm guard column, both with a 5 µm particle size. A binary solvent system was employed, where Solvent A consisted of 20 mM ammonium carbonate with 0.05% ammonium hydroxide, and Solvent B was pure acetonitrile. The column oven was maintained at 40 °C, and the autosampler tray was kept at 4 °C. Chromatography was performed at a flow rate of 0.200 mL/min with the following gradient program: 0–2 minutes at 80% Solvent B, followed by a linear gradient from 80% to 20% Solvent B between 2 and 17 minutes, a rapid return to 80% Solvent B from 17 to 17.1 minutes, and a hold at 80% Solvent B from 17.1 to 23 minutes. Samples, each injected at a volume of 5 µL, were randomized to minimize analytical variability. A pooled quality control (QC) sample, prepared by mixing equal volumes of all individual samples, was analyzed at regular intervals between test samples to ensure analytical consistency.

Metabolite quantification was conducted using a Vanquish Horizon UHPLC system connected to an Orbitrap Exploris 240 mass spectrometer (both Thermo Fisher Scientific), utilizing a heated electrospray ionization (HESI) source. The ionization parameters were set with spray voltages of +3.5LkV for positive mode and -2.8LkV for negative mode, an RF lens value of 70, a heated capillary temperature of 320 °C, and an auxiliary gas heater temperature of 280 °C. The sheath gas flow rate was set to 40, auxiliary gas to 15, and sweep gas was turned off. For MS1 scans, the mass range was established from m/z 70 to 900, employing a standard automatic gain control (AGC) target and an automatically set maximum injection time. Data acquisition of experimental samples was performed in full scan mode with polarity switching at an Orbitrap resolution of 120,000. The AcquireX Deep Scan workflow facilitated untargeted metabolite identification through an iterative data-dependent acquisition strategy, utilizing multiple pooled sample injections. The settings included a full scan resolution of 60,000, a fragmentation resolution of 30,000, and a fragmentation intensity threshold of 5.0L×L10³. Dynamic exclusion was enabled after a single occurrence, with a 10-second exclusion duration and a mass tolerance of 5Lppm. The isolation window was set to 1.2Lm/z, and normalized higher-energy collisional dissociation (HCD) collision energies were applied in stepped mode at 30, 50, and 150. Mild trapping was also enabled to enhance signal quality.

### Metabolomics data processing

Metabolite identification was performed using Compound Discoverer software (v3.2, Thermo Fisher Scientific). The identification criteria included a precursor ion m/z within 5Lppm of the theoretical mass based on the chemical formula, fragment ions within 5Lppm matching an internal spectral library of authentic compound standards analyzed using the same data-dependent MS² (ddMS²) method with a minimum match score of 70, and retention times within 5% of those of purified standards under identical chromatographic conditions. Peak area integration and chromatogram review were carried out using TraceFinder software (v5.0, Thermo Fisher Scientific).

### RNA extraction for transcriptomics

1x10^6 CD14+ monocytes were isolated as described above and optionally differentiated into M(IFNγ +LPS) or M(IL-4) macrophages as described. Following a 24-hour incubation with 100 ng/mL LPS or vehicle control, cells were washed twice in ice-cold PBS and pelleted by centrifugation (4°C, 400xg, 5min). RNA extraction and on-column DNA digestion was performed using the RNeasy Mini Kit (Qiagen) according to the manufacturer’s protocol.

### Illumina sequencing & data processing

Library preparation and sequencing were performed at the Genomics Facility Basel, Switzerland operated by the University of Basel and the Department of Biosystems Science and Engineering (D-BSSE), ETH Zurich. RNA integrity was assessed using an Agilent TapeStation system with RNA ScreenTape assays, yielding RNA integrity numbers (RINe) between 8.7 and 9.6 for all samples. Total RNA libraries were prepared using the TruSeq Stranded mRNA Library Preparation Kit (Illumina) from 200 ng total RNA, including poly(A) selection and strand-specific library construction. Libraries were sequenced on an Illumina NovaSeq 6000 platform using paired-end sequencing (∼2×100 bp) with dual 10-bp index reads on one lane of an S4 flow cell, generating approximately 25–30 million read pairs per sample.

The dataset was processed by the Bioinformatics Core Facility, Department of Biomedicine, University of Basel, Switzerland. Base calling and demultiplexing were performed using bcl2fastq (v2.20.0.422) allowing one mismatch in index reads. Raw sequencing reads were assessed for quality using FastQC, and summary quality metrics across samples were compiled using MultiQC. cDNA reads were aligned to the human genome (UCSC version hg38) with STAR (v2.7.10a) using default parameters except allowing up to 10 hits in the genome (outFilterMultimapNmax 10), reporting only one alignment per multi-mapping read (outSAMmultNmax 1), and filtering reads without evidence in the splice junction table (outFilterType “BySJout”). Alignments were sorted and indexed with samtools (v1.11). Gene-level read counts were generated using featureCounts from the Subread package (v2.0.1), counting read 5′ ends overlapping exons using the exon union model, with parameters -O, -M, --read2pos=5, --primary, -s 2, -p, -B. Gene annotation was based on Ensembl release 104. Only protein coding genes (*GENEBIOTYPE == “protein_coding”*) were used in downstream analyses.

### Cell culture & lysis for proteomics

Peripheral blood from HD and 8 RA patients was collected in EDTA tubes. CD14⁺ monocytes were isolated by magnetic bead separation and resuspended at 1 × 10L cells/mL. For each donor, cells were divided into three subsets corresponding to M0, M(IFNγ +LPS) and M(IL-4). Where necessary, higher cell numbers were allocated to M(IFNγ +LPS) and M(IL-4) cultures (e.g., from 8 × 10L CD14⁺ cells: 2 × 10L for M0, 3 × 10L for M1, and 3 × 10L for M2). Each subset was further split into two equal fractions to generate unstimulated and LPS-stimulated conditions and plated in 12-well plates. M0 cells were cultured without differentiation factors and stimulated, where indicated, with 100 ng/mL LPS for 24 h. For M(IFNγ +LPS) and M(IL-4) polarization, CD14⁺ monocytes were differentiated for 8–11 days with cytokine supplementation every 2–3 days (M(IFNγ +LPS): 20 ng/mL LPS plus IFN-γ; M(IL-4): 20 ng/mL IL-4). After differentiation, one well per condition was stimulated with 100 ng/mL LPS for 24 h, while the paired well remained unstimulated. Cells were harvested using pre-warmed EDTA/trypsin, neutralized with culture medium, and counted. A total of 5 × 10L cells per sample were washed twice with cold PBS (400 × g, 4 min, 4°C), snap-frozen in liquid nitrogen, and stored at −80°C. Upon completion of sample collection (6 conditions per donor; 96 samples in total), pellets were thawed, lysed in 100 µL lysis buffer, boiled at 95°C for 10 min, transferred to a 96-well plate, sealed, and stored at −20°C until further processing.

### Label-free LC-MS/MS proteomics

Proteins were purified and digested with the SP3 approach (50) using a Freedom Evo 100 liquid handling platform (Tecan Group Ltd., Männedorf, Switzerland). In brief, Speed BeadsTM (#45152105050250 and #65152105050250, GE Healthcare) were mixed 1:1, rinsed with water and diluted to the 8 μg/µL stock solution. Samples were adjusted to the final volume of 90 µL and 10 µL of the beads stock solution was added to them. Proteins were bound to the beads by addition of 100 µL of 100% acetonitrile to the samples, which were then incubated for 8 min at RT with a gentle agitation (200 rpm). After, samples were placed on a magnetic rack and incubated for 5 minutes. Supernatants were removed and discarded. The beads were washed twice with 160 µL of 70 % (v/v) ethanol and once with 160 of 100% acetonitrile. Samples were placed off the magnetic rack and 50 µL of digestion mix (10 ng/µL of trypsin in 50 mM triethylammonium bicarbonate) was added to them. Digestion was allowed to proceed for 12h at 37°C. After digestion samples were placed back on the magnetic and incubated for 5 minutes. Supernatants containing peptides were collected and dried under vacuum.

Dried peptides were resuspended in 0.1% aqueous formic acid and subjected to LC-MS/MS analysis using a Orbitrap Exploris 480 Mass Spectrometer (Thermo Fisher Scientific) fitted with a Vanquish Neo UHPLC System (Thermo Fisher Scientific,) and a custom-made column heater set to 60°C. Peptides were resolved using a reverse-phase high-performance liquid chromatography (RP-HPLC) column (75μm × 30cm), packed in-house with C18 resin (ReproSil-Pur 120 C18–AQ, 1.9 μm resin; Dr. Maisch GmbH), at a flow rate of 0.2 μL/min. The following gradient was used for peptide separation: from 2% Buffer B to 10% Buffer B over 5 min, to 35% Buffer B over 45 min, to 50% Buffer B over 10 min, to 95% Buffer B over 1 min followed by 10 min at 95% Buffer B. Buffer A was 0.1% FA and buffer B was 0.1% FA in 80% ACN.

The mass spectrometer was operated in data-independent acquisition (DIA) mode. Each MS1 scan was acquired at a resolution of 120,000 FWHM (at 200 m/z) and a scan mass range from 390 to 910 m/z. Maximum injection time was set to auto mode with a normalized AGC target set to 300%. Each survey scan was followed by a set of DIA scans acquired at a resolution of 15,000 FWHM. Precursor mass range was set to 400-900 m/z, the isolation windows size to 10 m/z and the HCD normalized collision energy (NCE) to 28%. For MS2 scans, maximum injection time was set to 22 ms with a normalized AGC target set to 3000%. 50 DIA overlapping windows were used to cover the mass range of interest.

### Proteomics data processing

The dataset was processed by the Proteomics Group at Functional Genomics Center Zurich, Switzerland. The acquired shotgun MS data were processed for identification and quantification using DIANN (51). Spectra were searched against a Uniprot Homo sapiens reference proteome (reviewed canonical version from 2023-03-30), concatenated to common protein contaminants. Post-translational modification on peptide N-termini and Lysine side chains and carbamidomethylation of cysteine were fixed modifications, while methionine oxidation was variable. Enzyme specificity was set to trypsin/P, allowing a minimal peptide length of 7 amino acids and a maximum of two missed cleavages.

From precursor ion abundances reported by DIA-NN, protein abundances were determined by first aggregating the precursor abundances to peptidoform abundances. Then Tukeys-Median Polish was employed to estimate protein abundances, followed by transformation using the variance stabilizing normalization (VSN) (52).

### Hierarchical clustering

For the transcriptomics dataset, lowly expressed genes (< 10 counts) were removed by expression-based filtering using *edgeR::filterByExpr()*, and library size differences were normalized with the trimmed mean of M-values (TMM) method using *edgeR::calcNormFactors*(). Variance-stabilized expression values were obtained via voom transformation using *limma::voom()*, yielding log2-CPM values suitable for downstream multivariate analyses. For the proteomics dataset, processed data as described above was directly used. For the metabolomics dataset, Probabilistic Quotient Normalization (PQN) (53) was performed to correct for variations in overall sample abundance, followed by feature-wise VSN transformation.

Pearson correlation as distance matrix and Ward agglomeration was used for general clustering analysis of all three omics datasets. For clustering heatmaps of protein abundances in Fig. 3, Euclidean distance and single-linkage agglomeration was used.

### Weighted Gene Co-expression Network Analysis (WGCNA)

Datasets were processed as described above for hierarchical clustering. Weighted Gene Co-expression Network Analysis (WGCNA) was subsequently performed on the voom-transformed expression matrix using the *WGCNA* R package. A signed network was constructed using biweight midcorrelation (*bicor*) to account for the directionality of gene-gene relationships. The soft-thresholding power for network construction was selected by evaluating scale-free topology model fit (R² threshold ≥ 0.80) across a range of candidate powers, with a power of 25 chosen for transcriptomics and 20 chosen for proteomics and metabolomics. Gene modules were identified using the *blockwiseModules()* function with the following parameters: signed topological overlap matrix (TOM), deep split sensitivity of 2, and a module merging height cutoff of 0.25. Minimum module size was set to 30 for transcriptomics, 15 for proteomics, and 3 for metabolomics.

Module eigengenes (MEs) representing the first principal component of gene expression within each module were calculated and used to characterize inter-module relationships. Intramodular connectivity (module membership, kME) was computed as the Pearson correlation between each gene’s expression profile and the module eigengenes. Module-trait correlations were calculated using *WGCNA::cor()* using all complete pairwise observations.

### Variance partitioning and differential analysis

Datasets were processed as described above for hierarchical clustering. To quantify the contribution of different biological factors to feature (transcript, protein or metabolite abundance) variability, variance partition analysis was performed on the normalized, transformed expression matrices using the *variancePartition* R package (54). All factors (donor, disease status, differentiation stage, LPS treatment) were modelled as random effects, and the proportion of variance attributable to each factor was estimated gene-wise using *fitExtractVarPartModel()*. To further resolve the sources of variance within the factor explaining the greatest median proportion of variance across genes, the dataset was stratified by the levels of that factor and variance partition analysis was repeated within each subset, excluding the stratifying factor from the model.

For differential analysis, a composite group factor was constructed representing the full interaction of disease status, differentiation stage, and LPS treatment, and a mixed-effects model was fit using the Dream framework (55), with donor modelled as a random effect and the interaction group as a fixed effect (intercept-free). Contrasts of interest were defined a priori and included pairwise comparisons of rheumatoid arthritis versus healthy donor macrophages within specific differentiation stage and treatment combinations, as well as comparisons across differentiation stages and LPS versus vehicle treatment within healthy donors. The model was fit with the pre-specified contrast matrix, and empirical Bayes moderation was applied via *eBayes* to stabilize variance estimates.

### Gene set enrichment analysis (GSEA) and Overrepresentation Analysis (ORA)

Gene set enrichment analysis (GSEA) was performed on transcriptomics and proteomics differential expression results using the *fgsea* R package. Six contrasts were selected for pathway analysis: rheumatoid arthritis versus healthy donor comparisons at each differentiation stage (M0, M(IFNγ +LPS), and M(IL-4).) under vehicle treatment, healthy donor M(IFNγ +LPS) and M(IL-4) versus M0 comparisons, and LPS versus vehicle treatment in healthy donor M0 macrophages. For each contrast, genes were ranked using the signed t-statistic from differential analysis results. A combined gene set collection comprised of KEGG pathways, Reactome pathways, Gene Ontology terms, MSigDB Hallmarks, MSigDB transcription factor targets, and MSigDB miRNA targets was used, with gene sets restricted to between 15 and 500 members.

Redundant significant pathways (adjusted p-value < 0.05) were first collapsed using *collapsePathways()*, applying a permutation p-value threshold of 0.005; if more than 30 main pathways remained after this step, collapsing was repeated with a stricter threshold of 0.0001 and 20,000 permutations to ensure a more parsimonious set of representative pathways. The resulting main pathways were further reduced using *rrvgo* (56) to consolidate semantically similar GO terms. Final results were annotated with leading-edge gene symbols and saved per contrast and ranking scheme.

The f*ora()* function was used for overrepresentation analysis (ORA) of member genes/proteins of WGCNA modules. The same combined gene set collection as for GSEA was used, with the union of all module-assigned genes serving as the background universe. Redundant significant pathways (adjusted p-value < 0.01) were collapsed per module using *collapsePathwaysORA()*, and the resulting representative pathways were further consolidated using rrvgo to reduce semantically similar GO terms. Results were annotated with overlapping gene symbols and combined into a single output table.

GSEA on MOFA weights was performed similarly as described above.

For pathway enrichment of metabolomics data, metabolite names were converted to KEGG IDs, and GSEA and ORA were performed as above, using KEGG pathways (restricted to between 5 to 100 members) as the ‘gene set’ collection.

### Integrated pathway-level analysis of metabolic pathways

Pathway-level integration of Integrated Molecular Pathway Level Analysis (IMPaLA) (57) was performed using the web interface at http://impala.molgen.mpg.de/ (accessed 9 October 2025). RA and HD group averages of normalized, variance-stabilized feature values were extracted from the differential analysis results (see above) as follows: the *AveExpr* column values were used for the HD group, while the sum of two columns *AveExpr* + *logFC* was used for the RA group. Metabolite KEGG Compound IDs, as well as gene ENSEMBL IDs for transcriptomics or UniProt IDs for proteomics, along with the two group averages were uploaded to the webserver for Wilcoxon enrichment analysis (WEA).

### Pseudosample pooling

Individual datasets were processed as described above for hierarchical clustering. For each omics layer, samples were grouped according to combinations of disease status, differentiation stage and treatment, generating composite pseudosamples representing group-level conditions. Linear modelling of group means was performed using the *limma* R package, whereby an ordinary least squares model with group indicators as design factors was constructed for each omics dataset using *lmFit()*, followed by empirical Bayes moderation *using eBayes()* to obtain stable estimates of condition-specific coefficients and standard errors. These group-level coefficients were used as pseudosample representations for integration across omics layers, while standard errors were used to perform feature filtering prior to downstream analyses including MOFA and Compass.

### Multi-Omics Factor Analysis (MOFA)

Multi-omics integration was performed using the MOFA2 framework (26) in R to identify latent factors capturing coordinated variation across transcriptomic, proteomic, and metabolomic datasets. Input matrices consisted of pseudosample-level group means derived from linear modelling of each omics dataset (see above), with features in rows and pseudosamples representing combinations of disease status, differentiation stage, and treatment in columns. Corresponding matrices of moderated standard errors were used to assess the reliability of each feature estimate. For quality control, relative standard errors (RSE; standard error divided by coefficient) were calculated, and feature–pseudosample values with RSE > 0.3 were set to missing. Features with missing values in at least half of the pseudosamples were subsequently removed. The resulting matrices were aligned to ensure identical pseudosample ordering across all omics layers and supplied as separate views to the MOFA model.

A MOFA model was initialized using default data and training parameters, with spike-and-slab priors enabled for factor weights to promote sparsity and improve interpretability of feature contributions. The model was trained using the MOFA2 implementation with variational Bayesian inference. The proportions of variance explained by each factor in each omics view, factor values for each pseudosample, and feature weights indicating the contribution of individual features (transcripts, proteins, and metabolites) to each latent factor were extracted from the model.

### Contraint-based metabolic modeling

Normalized feature abundance matrices were generated from transcriptomic and proteomic datasets for constraint-based metabolic modelling. Features were first filtered to remove low-confidence measurements: for transcriptomics, lowly expressed genes were filtered out using *filterByExpr()* from the *edgeR* package, while for proteomics, proteins detected in fewer than 70% of samples were excluded. Sample-wise normalization was then applied to account for differences in library size or global signal intensity: transcriptomic counts were normalized using the trimmed mean of M-values (TMM) method and expressed as counts per million (CPM), while proteomic abundances were normalized using probabilistic quotient normalization (PQN).

For integrative transcriptomics- and proteomics-based metabolic modelling, group-averaged pseudosample matrices were calculated from the normalized abundance matrices, using unique combinations of disease status, differentiation stage and treatment for pseudosample definition as performed above for MOFA. Feature names in the two matrices were first harmonized using HGNC gene symbols as the common identifier. The two matrices were then merged, following a hybrid integration strategy in which proteomic values were preferentially retained when available, while transcriptomic values were used to fill missing entries. This produced a single gene-by-condition matrix representing integrated abundance estimates for all experimental pseudosamples, which was exported for downstream metabolic modeling.

Metabolic activity inference was subsequently performed using Compass (v1.0.0) (27). The combined gene expression matrix was supplied as input, and reaction activity scores were inferred using the human metabolic reconstruction RECON2 model under default media conditions. Compass maps gene expression values to metabolic reactions through gene–protein–reaction rules and computes reaction penalties reflecting the likelihood of reaction usage given the observed expression levels. Flux potentials were then estimated for each reaction and sample using a linear programming framework, with diffusion-based penalty smoothing applied using a k-nearest neighbor graph (k = 30).

Compass reaction penalties were converted into reaction activity scores, using a negative logarithmic transformation (score = -log(penalty + 1)) such that higher values indicate greater predicted reaction activity, followed by rescaling so that the minimum value for each reaction across samples was set to zero. Reactions showing minimal variability across samples were excluded, and analyses were restricted to curated core metabolic reactions with high or unassigned confidence annotations, valid Enzyme Commission numbers, and appropriate cellular localization. Differential analysis was performed on the resulting reaction scores using linear mixed-effects modeling implemented by *dream()* of the *variancePartition* R package, as described above.

### Metabolomics validation cohort data acquisition and analysis

For validation of the metabolomic signature in RA monocytes, CD14+ monocytes were obtained from an independent cohort of RA patients and healthy donors for metabolomics analysis.

Study ethics and participants, as well as isolation of CD14+ monocytes were as described above. Following CD14+ isolation, cells were counted using a Countess 3 FL Automated Cell Counter (Thermofisher) and 4 million monocytes were pelleted by centrifugation, washed by resuspending in room temperature PBS and re-pelleted by centrifugation, then snap frozen in liquid nitrogen and stored at -80°C until cohort collection was complete. Metabolite extraction was then performed by adding 500µL of ice-cold mixed solvent (40% LC-MS grade methanol, 40% LC-MS grade acetonitrile, 20% LC-MS grade water, and containing 1 µM 1,4-Piperazinediethanesulfonic acid (PIPES) as internal standard), followed by vortexing and centrifugation at 4°C, 10,000 g for 10 min. The upper 80% of the supernatant was carefully collected, dried under a stream of nitrogen gas, then stored at -80 C until LC-MS/MS analysis. The protein pellet was dried, dissolved in 0.5 M potassium hydroxide and quantified using Pierce BCA Protein Assay Kit (Thermofisher) as a measure of input cell amount for normalization purposes.

Dried metabolite extracts were reconstituted in 80% LC-MS grade water, 20% LC-MS grade acetonitrile for analysis. Metabolites were separated chromatographically on a Thermo Vanquish Horizon UHPLC system using a Waters ACQUITY HSS T3 column (2.1L×L150Lmm) with a 1.8 µm particle size. A binary solvent system was employed, where Solvent A consisted of water with 0.1% formic acid and Solvent B consisted of acetonitrile with 0.1% formic acid. The column oven was maintained at 40 °C, and the autosampler tray was kept at 4 °C. Sample injection volume was 3 µL. Chromatography was performed with the following gradient program: t = 0 min, 0.250 mL/min flow rate, 0% B; t = 1 min, 0.300 mL/min flow rate, 0% B; t = 6 min, 0.300 mL/min flow rate, 20% B; t = 11 min, 0.300 mL/min flow rate, 100% B; t = 14 min, 0.450 mL/min flow rate, 100% B; t = 14.1 min, 0.250 mL/min flow rate, 0% B; t = 16 min, 0.250 mL/min flow rate. The chromatograph was coupled to a Thermo Q Exactive HF mass spectrometer via a heated electrospray ionization (HESI) source. The ionization parameters were set with spray voltages of +3.8LkV for positive mode and -3.5LkV for negative mode, an

S-lens RF level of 50, and heated capillary temperature of 300 °C. The sheath gas flow rate was set to 30 and auxiliary gas to 10. Data acquisition of experimental samples was performed in full scan mode with polarity switching with mass range from m/z 70 to 1050, Orbitrap resolution of 60,000, automatic gain control (AGC) target of 3e6 and maximum injection time of 100 ms. Pooled QC samples were injected twice every 8 experimental samples for Data Dependent Acquisition (DDA), once each with positive and negative ionization modes respectively. The MS1 scan settings included mass range from m/z 70 to 1050, resolution of 60,000, AGC target of 1e6, and maximum injection time of 100 ms. MS2 acquisition settings included TopN of 5, intensity threshold of 1.6e5, isolation window of 1.0 m/z, dynamic exclusion window of 4.0 s, nominal collision energies (NCE) of 10, 20 and 40, mass range from m/z 200 to 2000, resolution of 15,000, AGC target of 1e5, minimum AGC target of 8e3, and maximum injection time of 50 ms.

Raw data files were converted to mzML format using ProteoWizard msConvert (58). Peak detection, alignment and annotation were performed using Mzmine 4 (version 4.8.30). MS2 spectral libraries used for peak annotation were from MassBank of North America (All LC-MS/MS Orbitrap, downloaded on 31 July 2025) and MassBank EU (NIST format, downloaded on 17 June 2025). Exported peak areas were normalized to the PIPES internal standard peak, then further normalized for differences in input sample amount using the BCA quantified protein values.

### Transcriptomics validation dataset analysis

For validation of transcriptomic signatures of RA vs HD monocytes, we utilized publicly available transcriptomics data acquired on isolated CD14+ monocytes from the RA-MAP consortium (25). The processed data matrix file was downloaded from Gene Expression Omnibus (GEO) dataset GSE97810. After filtering for CD14+ samples (*tissue* == “PBMC_CD14”), RA samples were defined as having either *x.clinical.data.00.baseline.acpa.acpa.positive.00bl: Yes* or *x.clinical.data.00.baseline.rheumatoid.factor.rheum.factor.positive.00bl: Yes* under the *!Sample_characteristics_ch1* metadata column, corresponding to samples that were tested and found positive for either anti-citrullinated protein antibody or rheumatoid factor at baseline. Differential gene expression analysis was performed on the processed microarray intensity values for 188 RA versus 23 control samples using *limma* (59). GSEA was performed using the moderated t statistic from differential analysis, using gene set collections as described above.

## ACKNOWLEDGEMENTS

RNA-seq library preparation and sequencing were performed at the Genomics Facility Basel, operated by the University of Basel and the Department of Biosystems Science and Engineering, ETH Zurich. External validation of the metabolomics dataset was made possible with support from Kristina Bürki at University Hospital Zurich for coordinating patient sample acquisition, and Dr. Alaa Othman, Dr. Martina Zanella and Dr. Annika Jagels from the Functional Genomics Center Zurich for support in metabolomics data acquisition.

## AUTHOR CONTRIBUTIONS

**Shao Thing Teoh**: Writing – Original Draft Preparation (lead); Data Curation (supporting); Formal Analysis (lead); Methodology (equal); Validation (equal); Visualization (lead). **Silja Malkewitz**: Investigation (equal); Validation (equal). **Cristian Iperi**: Data Curation (equal), Formal Analysis (supporting); Validation (equal). **Celia Makowiec**: Data Curation (equal); Investigation (equal). **Asimina Kakale**: Investigation (supporting); Validation (supporting). **Seyram Maureen Duphey**: Data Curation (supporting); Investigation (supporting). **Anastasiya Börsch**: Data Curation (equal); Formal Analysis (supporting); Methodology (equal). **Katarzyna Buczak**: Data Curation (equal); Investigation (equal). **Witold Wolski**: Data Curation (equal); Formal Analysis (supporting); Methodology (equal). **Ming Yang**: Data Curation (equal); Formal Analysis (supporting); Investigation (equal). **Christian Frezza**: Project Administration (supporting); Resources (supporting); Supervision (supporting). **Caroline Ospelt**: Project Administration (supporting); Resources (supporting). **Oliver Distler**: Project Administration (supporting); Resources (supporting). **Diego Kyburz**: Project Administration (supporting); Resources (supporting). Bojana Mue**ller-Durovic**: Conceptualization (lead); Funding Acquisition (lead); Methodology (equal); Project Administration (lead); Resources (equal); Supervision (equal); Writing – Review & Editing (lead).

## DATA AVAILABILITY

Metabolomics data generated during this study were deposited at MetaboLights under the study accessions MTBLS14227 and MTBLS14228. The transcriptomics count data were deposited at Gene Expression Omnibus (GEO) as Dataset GSE328878. The proteomics data were deposited to the ProteomeXchange Consortium via the PRIDE (60) partner repository with the dataset identifier PXD076687.

---

**Figure S1.**
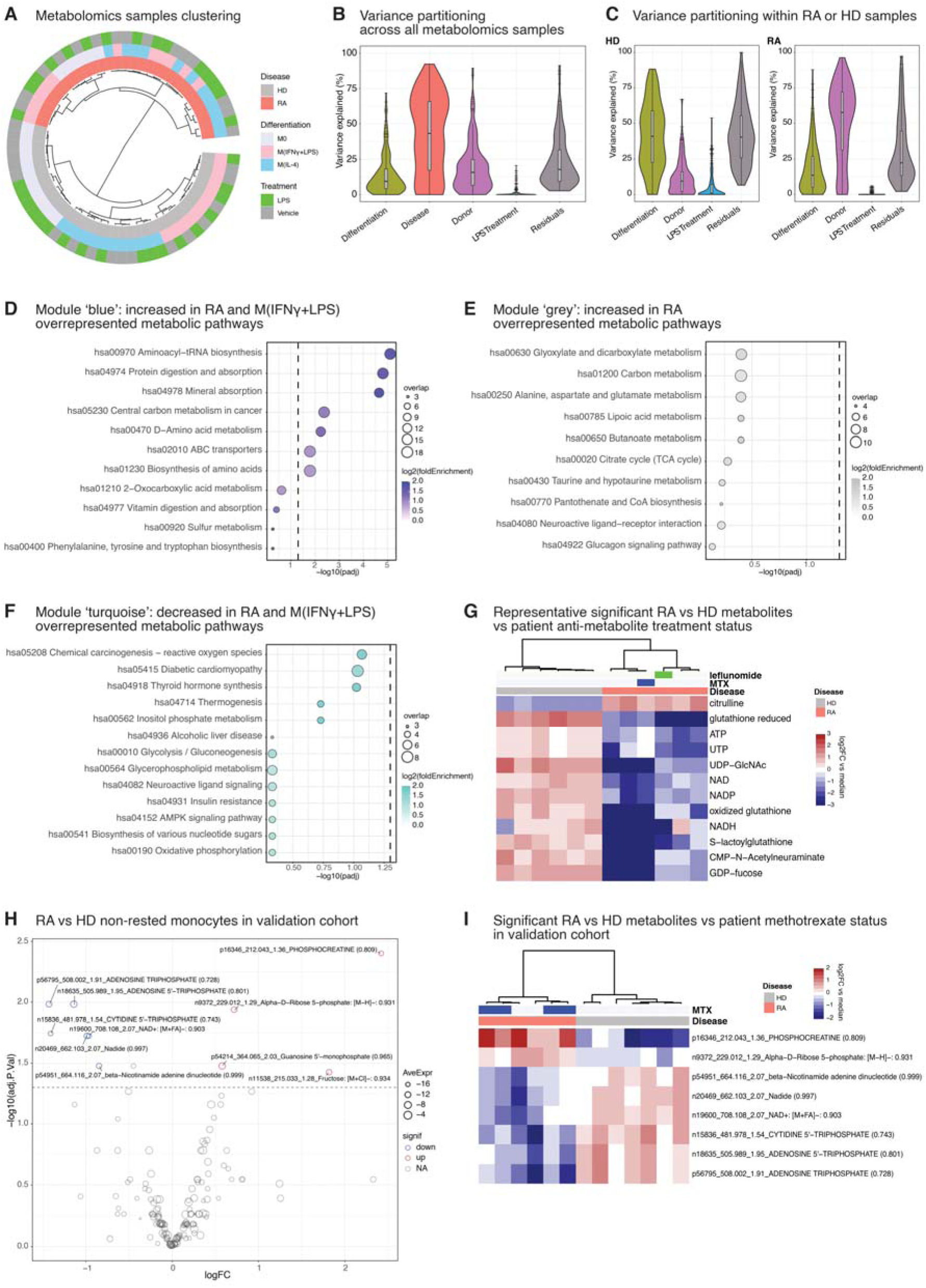
RA disease status dominates metabolomic variance. **(A)** Circular dendrogram showing hierarchical clustering of metabolomics profiles. Tree leaves represent samples and are annotated with sample disease status, differentiation state and LPS treatment status. The samples are clearly separated into two major clusters according to RA disease status, while within the HD samples the samples are further subclustered by differentiation state. **(B)** Violin plot showing metabolomics data variance explained by each sample factor – differentiation state, disease status, donor individuality, LPS treatment – or unexplained residuals. The violin plot shapes reflect the distribution of explained variance across all genes per sample factor, with nested boxes indicating the 1^st^ quartile, median and 3^rd^ quartile, while horizontal marks indicate individual genes with explained variance larger than 1.5 × interquartile distance above the 3^rd^ quantile (Tukey’s rule). **(C)** Violin plots showing metabolomics data variance explained by each sample factor within RA and HD sample subsets. **(D–F)** Overrepresentation analysis (ORA) of ‘blue’, ‘grey’, and ‘turquoise’ WGCNA modules, showing metabolic pathways enriched by the metabolites belonging to each module. Point colors and sizes indicate fold enrichment and number of overlap genes respectively. Vertical dashed lines indicate the adjusted p-value = 0.05 threshold. Top 10 metabolite pathways with smallest adjusted p-value are shown per plot. **(G)** Heatmap of representative significant RA vs HD metabolites in M0 monocytes confirming absence of correlation between methotrexate or leflunomide treatment and the RA-associated metabolic signature. Metabolite abundances are expressed as log2 fold-change over median peak abundance of the respective metabolite within the dataset. **(H)** Volcano plot showing differential metabolite abundance between RA and HD monocytes in a validation cohort (6 RA patients vs 7 healthy donors). Samples in this cohort were harvested directly for metabolomics analysis following CD14+ isolation, without overnight resting or differentiation. The horizontal axis indicates log2 fold change, while vertical axis indicates adjusted p-value of RA vs HD differences. Average metabolite peak areas are indicated by data point sizes. Significant metabolites (adjusted p-value < 0.05, absolute log2FC > 1) are color coded red or blue depending on log2FC direction and are labeled to aid interpretation. The results from this cohort confirm decreased nucleotides and NAD in RA monocytes. **(I)** Heatmap of significant RA vs HD metabolites in the validation cohort confirming absence of correlation between methotrexate treatment and the metabolic signature of RA monocytes. Metabolite abundances are expressed as log2 fold-change over median abundance of the respective metabolite within the dataset.

**Figure S2.**
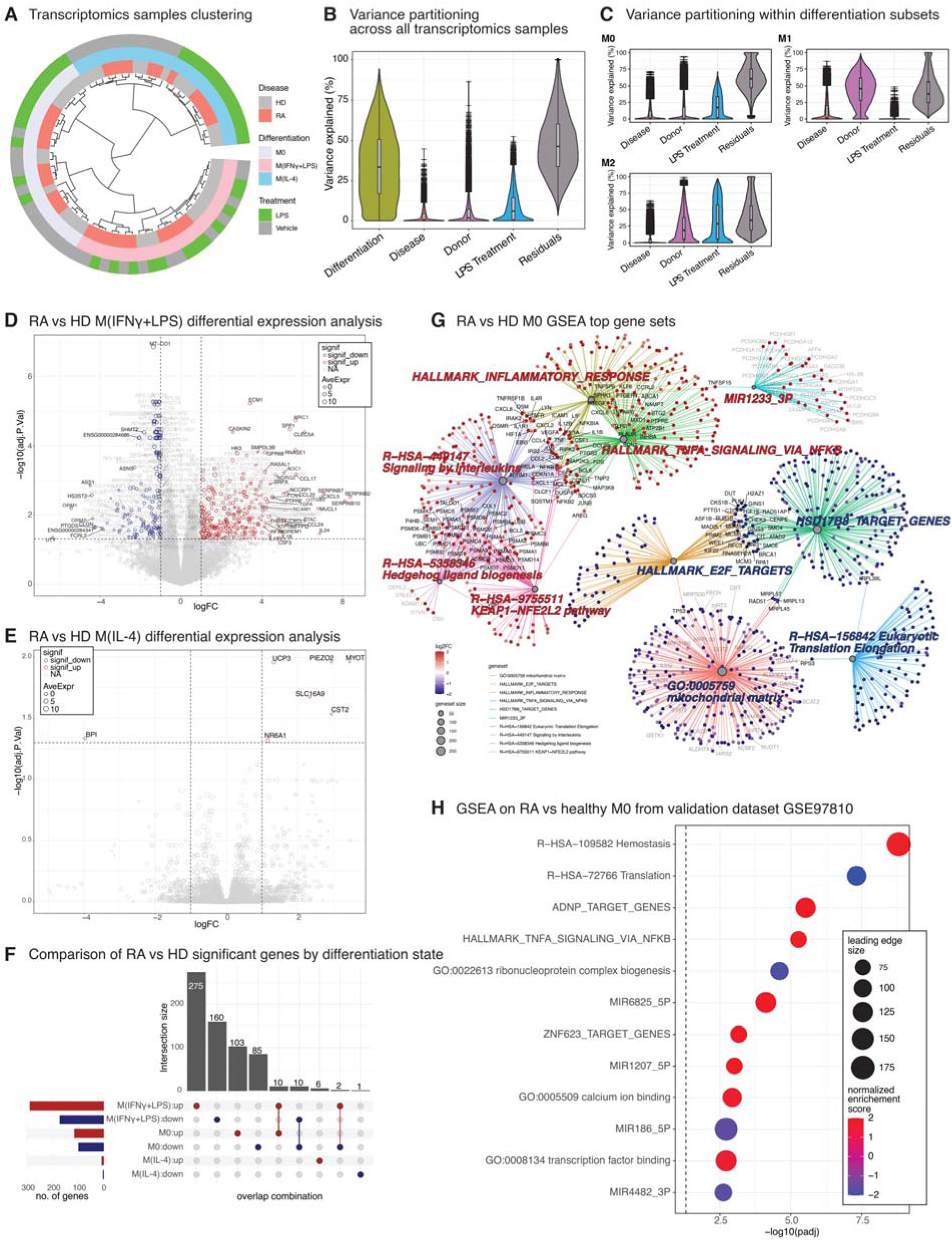
Transcriptomic variance is driven primarily by differentiation state. **(A)** Circular dendrogram showing hierarchical clustering of transcriptomics profiles. Tree leaves represent samples and are annotated with sample disease status, differentiation state and LPS treatment status. The samples are clearly separated into three major clusters reflecting differentiation state, while within M0 and M(IL-4) clusters the samples are further separated by LPS treatment status. **(B)** Violin plot showing transcriptomics data variance explained by each sample factor – differentiation state, disease status, donor individuality, LPS treatment – or unexplained residuals. The violin plot shapes reflect the distribution of explained variance across all genes per sample factor, with nested boxes indicating the 1^st^ quartile, median and 3^rd^ quartile, while horizontal marks indicate individual genes with explained variance larger than 1.5 × interquartile distance above the 3^rd^ quantile (Tukey’s rule). **(C)** Violin plots showing transcriptomics data variance explained by each sample factor within M0, M(IFNγ+LPS), and M(IL-4) subsets. **(D–E)** Volcano plots of RA vs HD differential gene expression within **(D)** M(IFNγ+LPS) differentiated, vehicle-treated samples, and **(E)** M(IL-4) differentiated, vehicle-treated samples. Each analysis compared 6 RA patients vs 6 healthy donors. The horizontal axis indicates log2 fold change, while vertical axis indicates adjusted p-value of RA vs HD differences. Point sizes indicate average gene expression level (normalized counts). Significantly up- or downregulated genes (adjusted p-value < 0.05, absolute log2FC > 1) are color-coded red or blue respectively. Selected genes are labeled to aid interpretation. **(F)** UpSet plot summarizing numbers of RA vs HD differentially expressed genes (adjusted p-value < 0.05, absolute log2FC > 1) across M0, M(IFNγ+LPS), and M(IL-4) differentiation states. Each row in the lower-right matrix represents a ‘category’ comprising a distinct differentiation state + direction of differential expression pairing, while columns represent overlaps between these categories. The bar plot on the left shows the number of genes in each category, while the bar plot above indicates number of genes overlapping between categories and reflects the limited overlap of differentially expressed genes across differentiation states. **(G)** Network representation of overlap between significantly enriched gene sets from transcriptomics RA vs HD GSEA. Top 10 gene sets with smallest adjusted p-value are shown. Gene sets are represented by grey filled circles with sizes according to number of leading edge genes. Leading edge genes are represented by nodes color-coded according to RA vs HD log2FC. Selected genes are labeled to aid interpretation. **(H)** GSEA using moderated t statistics from RA vs HD gene expression differential analysis of CD14+ monocytes from the RA-MAP validation cohort. The analysis included 188 RA patients vs 23 healthy controls. Point colors indicate normalized enrichment score (NES) whereby positive and negative scores indicate up- and downregulation in RA respectively, while point sizes indicate number of leading edge genes. The vertical dashed line indicates the adjusted p-value = 0.05 threshold. Top 12 gene sets with adjusted p-value < 0.05 are shown.

**Figure S3.**
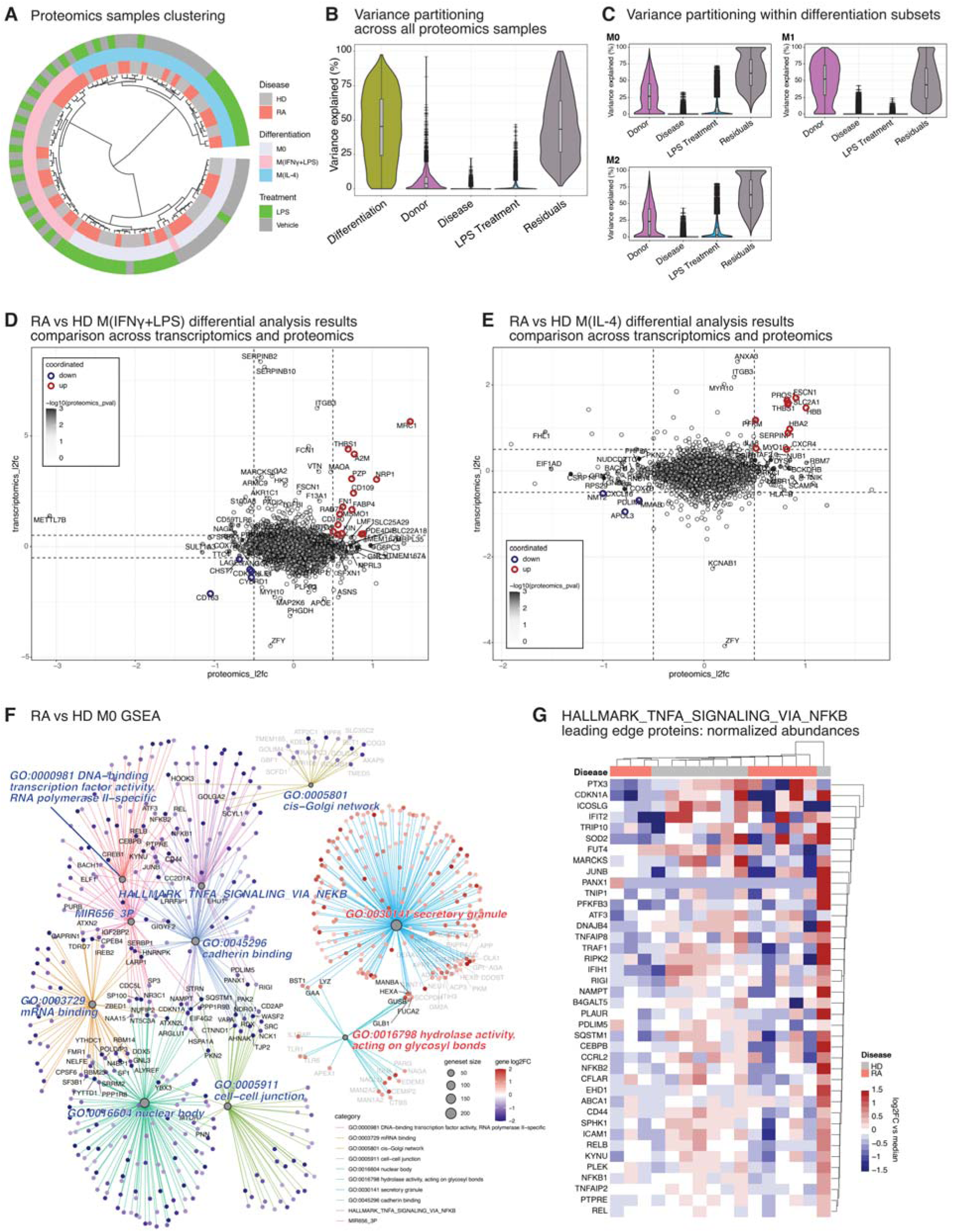
Proteomics analysis details and pathway-level alterations. **(A)** Circular dendrogram showing hierarchical clustering of transcriptomics proteomics profiles. Tree leaves represent samples and are annotated with sample disease status, differentiation state and LPS treatment status. The samples are clearly separated into three major clusters reflecting differentiation state, while within M0 and M(IL-4) clusters the samples are further separated by LPS treatment status. **(B)** Violin plot showing proteomics data variance explained by each sample factor – differentiation state, disease status, donor individuality, LPS treatment – or unexplained residuals. The violin plot shapes reflect the distribution of explained variance across all genes per sample factor, with nested boxes indicating the 1^st^ quartile, median and 3^rd^ quartile, while horizontal marks indicate individual genes with explained variance larger than 1.5 × interquartile distance above the 3^rd^ quantile (Tukey’s rule). **(C)** Violin plots showing proteomics data variance explained by each sample factor within M0, M(IFNγ+LPS), and M(IL-4) subsets. **(D–E)** Scatter plots comparing RA vs HD log2 fold-changes from transcriptomic and proteomic differential analyses in **(D)** M(IFNγ+LPS) differentiated, vehicle-treated samples, and **(E)** M(IL-4) differentiated, vehicle-treated samples. The transcriptomics differential analysis compared 6 RA vs 6 HD, while the proteomics analysis compared 8 RA vs 8 HD samples; all samples were biologically independent. The vertical and horizontal axes represent transcriptomics and proteomics log2FCs respectively. Data points are additionally shaded according to proteomics RA vs HD p-value. Genes up- or downregulated consistently in both transcriptomics and proteomics (above absolute log2FC threshold of 0.5) are outlined red or blue, respectively. Specific genes are labeled to aid interpretation. **(F)** Network representation of overlap between significantly enriched gene sets from proteomics RA vs HD GSEA. Top 10 gene sets with smallest adjusted p-value are shown. Gene sets are represented by grey filled circles with sizes according to number of leading edge proteins. Leading edge proteins are represented by nodes color-coded according to RA vs HD log2FC. Selected proteins are labeled with corresponding HGNC symbols to aid interpretation. **(G)** Clustered heatmap of leading-edge proteins from the *HALLMARK_TNFA_SIGNALING_VIA_NFKB* gene set demonstrating coordinated RA-associated abundance shifts. Protein abundances are expressed as log2 fold-change over median abundance of the respective protein within the dataset.

**Figure S4.**
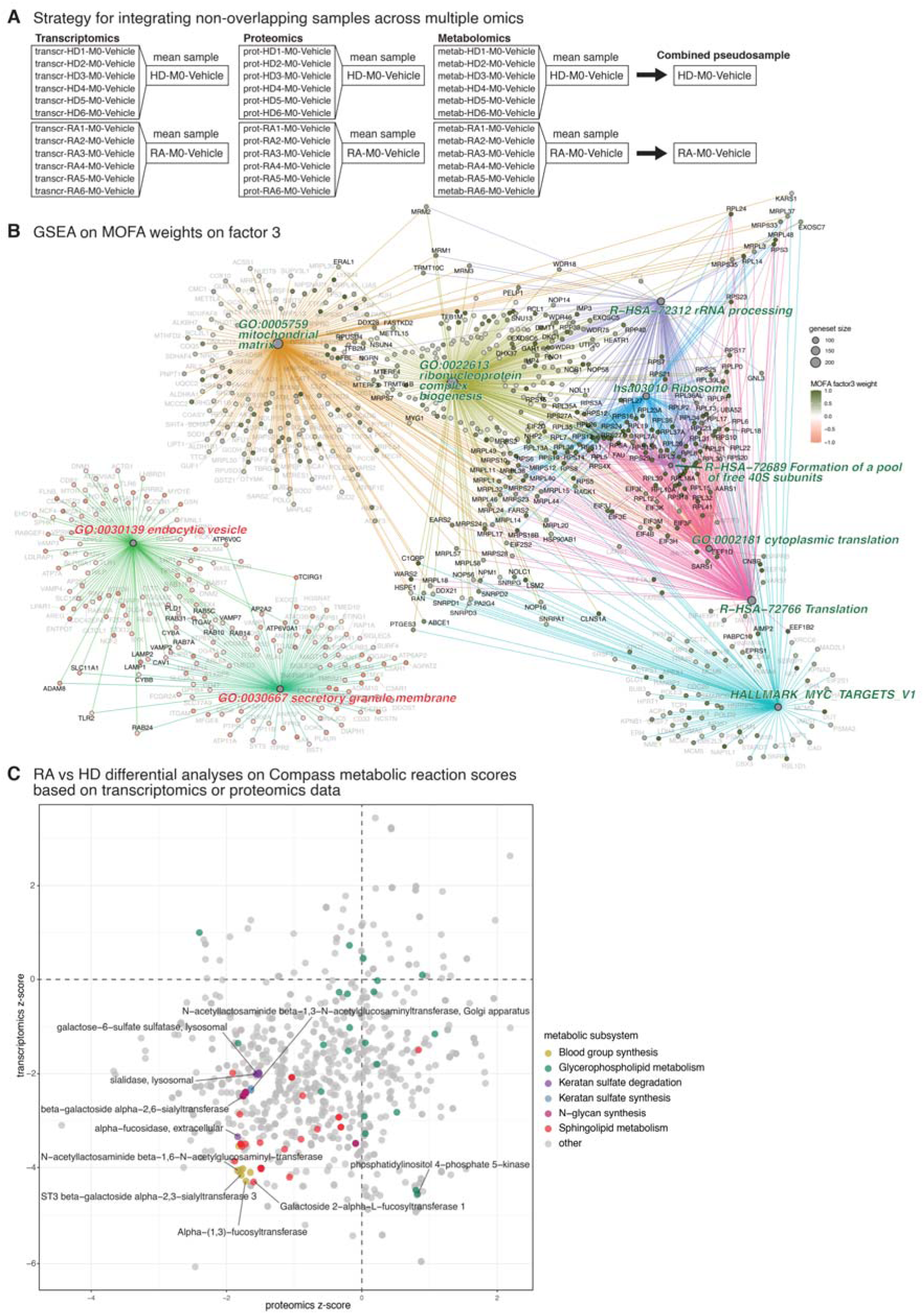
Multi-omics integration and metabolic modeling robustness analyses. **(B)** Schematic illustration of pseudosample pooling strategy for MOFA integration. Within each omics dataset, samples belonging to the same combination of disease status, differentiation state and LPS treatment status are pooled to obtain a single representative profile, the ‘pseudosample’ (see Methods). Each dataset yields 12 pseudosamples representing distinct combinations of sample factors. These pseudosamples are common across all three omics datasets, allowing downstream integrative analyses requiring matched samples (e.g. MOFA). **(C))** Network representation of overlap between significantly enriched gene sets from GSEA on MOFA Factor 3 weights. Top 10 gene sets with smallest adjusted p-value are shown. Gene sets are represented by grey filled circles with sizes according to number of leading edge genes. Leading edge genesare represented by nodes color-coded according to RA vs HD log2FC (salmon color for low weights corresponding to higher abundance in RA; olive green for high weights corresponding to higher abundance in HD). Selected genes are labeled with corresponding HGNC symbols to aid interpretation. **(D)** Scatter plot comparing RA vs HD differential reaction scores calculated from transcriptomics and proteomics profiles of untreated M0 monocytes. Reaction scores were first calculated from transcriptomics or proteomics profiles using Compass, then RA vs HD differential analysis was performed separately for each set of reaction scores. The transcriptomics differential analysis compared 6 RA vs 6 HD, while the proteomics analysis compared 8 RA vs 8 HD samples; all samples were biologically independent. The vertical and horizontal axes represent z-scores from transcriptomics- and proteomics-based differential analyses respectively. Metabolic subsystems related to cellular glycosylation and found in the cluster of reactions shown in Fig. 4F are highlighted here.

## Notes

**Funding:** This research was supported by SNSF Ambizione (PZ00P3_216467) (BMD), Uniscientia Stiftung (222-2024) (STT), Goldschmidt-Jacobson Foundation Basel (BMD), Alexander von Humboldt Foundation (CF, MY) and the Deutsche Forschungsgemeinschaft (CF, MY).

**Competing interests:** The authors declare no competing interests.

### Competing Interest Statement

The authors have declared no competing interest.

